# HAdV-5 infection dysregulates cysteine, purine, and unsaturated fatty acid metabolism in fibroblasts

**DOI:** 10.1101/2024.06.03.597117

**Authors:** Bailey-J C. Sanchez, Rudy M. Ortiz, Juris A. Grasis

## Abstract

Viral infections can cause cellular dysregulation of metabolic reactions. Viruses alter host metabolism to meet their replication needs. The impact of viruses on specific metabolic pathways is not well understood, even for a well-studied virus-like human adenovirus. Adenoviral infection is known to affect cellular glycolysis and respiration, however, global effects on cellular metabolic pathways in response to adenoviral infection are lacking, particularly in normally quiescent structural cells, such as fibroblasts. Further, few studies have employed an untargeted approach with an emphasis on viral dosage and duration of infection. To address this, we employed untargeted metabolomics to quantify the dynamic metabolic shifts in fibroblasts infected with human adenovirus serotype 5 (HAdV-5) at three dosages (0.5, 1.0, and 2.0 multiplicity of infection [MOI]) and across four time points (6, 12, 24, and 36 h post-infection [HPI]). The greatest differences in individual metabolites were observed at 6– and 12-hours post-infection. In addition to its effects on glycolysis and respiration, adenoviral infection downregulated cysteine and unsaturated fatty acid metabolism, while upregulated purine metabolism. These results reveal the specific metabolic pathways that are perturbed by adenoviral infection and the associated dynamic shifts in metabolism, suggesting that viral infections alter energetics via profound changes in protein, lipid, and nucleic acid metabolism. The results revealed previously unconsidered metabolic pathways disrupted by HAdV-5 that can alter cells, even in non-excitable structural cells, such as fibroblasts.

**Importance:** Human adenoviruses overtake the DNA replication machinery of the infected host, rewiring mitotic events and leading to effects on cellular respiration and glycolysis. Fibroblast lineages are normally quiescent cells that display a repertoire of responses to certain agonists. While metabolism often begins with glucose breakdown in the form of aerobic glycolysis, additional pathways are important for the overall functioning of the cell. Data on shifts in the metabolism of fibroblast cells in response to human adenoviral infection are lacking. We used an untargeted metabolomic approach to better understand the dynamic metabolic changes in human kidney cells in response to three viral dosages across four time points post infection. Profound shifts were observed for the cysteine, purine, and unsaturated fatty acid metabolites. This analysis provides a global perspective and highlights previously underappreciated aspects of how human adenoviruses alter host metabolism.

## Introduction

Cellular metabolism encompasses the chemical reactions necessary for livelihood. These reactions, often presumed as linear sequences, are multidirectional, with alterations in one pathway potentially affecting multiple pathways (1). Different cell types in the body yield a diverse array of functions, and one cell type may possess a different metabolic landscape compared to another. Fibroblasts, which are normally quiescent cells derived from the mesoderm, are versatile in their functionality and adaptability.

Metabolically, fibroblast activation is coupled with reprogramming, including the upregulation of aerobic glycolysis and glutaminolysis (2,3). Fibroblast metabolism has also been targeted for therapeutic purposes. For instance, a recent study showed that inhibition of the glycolytic enzyme 6-phosphofructo-2-kinase/fructose-2,6-biphosphatase 3 (PFKFB3) ameliorated pulmonary fibrosis in mice (3). When exposed to agonists, including viruses, fibroblasts respond with massive growth and proliferation, leading to the production of extracellular matrix material (4,5). Similarly, fibroblast metabolism is likely to be altered by viral infections.

Given that viruses are obligate intracellular parasites that are only capable of replicating within an infected host cell, they must co-opt multiple host pathways to enhance replication, including cellular metabolism. This cellular takeover provides the virus with the substrates and energy required for successful replication (6–8). While it has been understood that viruses alter host metabolism, only in the past decade have studies extensively examined these interactions, largely because of the development of metabolomic platforms, metabolite libraries, and analytical tools (9–11). Despite these advancements, multiple studies have employed targeted approaches, concerned with alterations to glycolysis and respiration, with less regard for additional components of overall cellular metabolism.

Human adenoviruses (HAdVs) are double-stranded DNA viruses capable of infecting the GI tract, kidneys, ocular regions, and respiratory tract (12). Currently, there are approximately 110 specific types of HAdVs across seven species termed A-G, each possessing extensive tissue tropism, which can be attributed to the presence of specific protein targets, including coxsackie and adenovirus receptor (CAR), and membrane cofactor proteins such as CD46 (13–15). HAdV pathogenicity may affect host metabolism, even at the level of a single protein expressed by the virus. The E4ORF1 isoform has been shown to disrupt glycolytic and glutamine metabolism *in vitro* (8,16). Additionally, the 13S E1A isoform, expressed in both wild-type and mutant forms, upregulates glycolytic genes and downregulates genes involved in cellular respiration (17). However, *in vitro* studies employing metabolomic techniques that capture the global metabolic profile of HAdV infections are lacking. Methods including ^1^H-NMR spectroscopy have been used to understand the subversion of cell metabolism from adenoviral infection, although the number of identified metabolites using this technique is less than that using mass spectroscopy techniques (18,19). Furthermore, studies examining the relationship between HAdV and host metabolism, with an emphasis on both the dose and duration of infection are also lacking.

Human adenovirus serotype 5 (HAdV-5) early transcript expression begins 3-4 hours after infection, notably E1A and E2A gene expression, with late-phase expression beginning 11-13 hours after infection (20–22). As the infection persists, viral transcripts continue to be expressed for up to 36 h, although early transcript plateauing has been reported beginning at 24 h. Studies conducted on adenoviral usurping of metabolism show an increased viral titer (10– and 50-fold compared to the cell number at the time of infection). However, it is important to assess the virulence of adenoviruses at lower titers. Therefore, we assessed the effects of HAdV-5 infection on cellular metabolism using gas chromatography (GC) coupled with time-of-flight mass spectrometry (TOF-MS), combining the infection period with multiple dosages. We hypothesized that the disruption of cellular metabolites would be proportional to the increase in viral dosage, with recovery after 24 h. Our study design allowed us to characterize the dynamic metabolic responses of human fibroblasts to adenoviral infection, yielding insights into the viral-induced shifts in metabolism from the early-onset infection to late-phase, cell death.

## Materials and Methods

### 2.1: Cell Culture

Human embryonic kidney (HEK293) cells were obtained from the American Type Culture Collection (CRL-1573). Cells were incubated in Dulbecco’s modified Eagle’s medium (DMEM) containing low glucose, phenol red, and pyruvate (Fisher Scientific) and mixed with 10% fetal bovine serum (Fisher Scientific) and 1% penicillin-streptomycin (Fisher Scientific). Incubator settings were maintained at 37.0 °C and %CO_2_ at 5.0. Cells were passaged every 2-3 days and split at 80–90% confluency, in which the expended media was discarded, and the wells were washed with serum-free DMEM and treated with 0.25% trypsin-EDTA (Fisher Scientific). After it was confirmed that the cells had lifted from the flask, they were transferred to a new T-25 flask at a 1:3 ratio, with fresh complete growth media added to adequately cover the flask. Cell counts and viability were determined using trypan blue staining and a hemocytometer. Cells used in the experiments were not used after passage 20.

### 2.2: HAdV-5 Propagation

HAdV-5 was obtained from the American Type Culture Collection (VR-1516). To acquire a highly concentrated viral stock, HEK293 and human osteosarcoma epithelial cells (U-2 OS; American Type Culture Collection HTB-96) were used for viral propagation. For viral stock propagation, after 3 days of HAdV-5 infection in a T-75 flask, cells were trypsinized, media was added, and the cells were transferred to a centrifuge tube. The cells were then pelleted, and three freeze/thaw cycles were performed, in which the samples were placed in liquid nitrogen for 30 s for each cycle. The cells were then transferred to a 15 mL, 50 kDa ultracentrifuge filter unit (Fisher Scientific). The filter units were spun at 2000 rpm for 20 min in a swinging-rotor centrifuge unit (Beckman Coulter). The retained product containing the virus was then used to infect new host cells. This was repeated three more times, and the cells were stored at –80.0 °C.

### 2.3: Reverse Transcription Quantitative PCR (RT-qPCR)

To confirm viral infection and quantify viral titers, the infected cells were subjected to RT-qPCR. Media harboring infected cells were removed, cells were washed with serum-free media, and cells were treated with guanidine isothiocyanate lysis buffer (TRIzol, ThermoFisher) (23). cDNA synthesis was performed according to the manufacturer’s guidelines using the SuperScript III Reverse Transcriptase kit (ThermoFisher) (24). The adenoviral genes of interest used for this analysis were E2A and E4. Primers were designed *de novo* using Primer-BLAST (Supplementary Table 1). Primers were designed to span all regions indicated by the HuAdV5-PanE2A and HuAdV5-PanE4 identifiers, which have been previously validated in SYBR-based dye qPCR assays (25). GAPDH was used as a control gene. qPCR reactions to confirm viral infection were performed using an Eco Real-Time PCR System (Illumina). Viral titer was achieved by using a modified SYBR-based dye assay protocol featuring the HuAdV5-PanE1A primer set (26).

### 2.4: Intracellular Metabolite Profiling

To perform viral infections for metabolomic analyses, the cells were passaged into 6-well plates. The cell density at the time of infection was 1.0 × 10^6^ cells/mL. The cells were infected for 6, 12, 24, and 36 h. Non-infected cells cultured for 24 h were used as the negative controls. Cells were subjected to the following viral dosages: 0.5, 1.0, and 2.0 multiplicity of infection (MOI), and six replicates were used for each experimental condition. Cellular pellets were collected at each time point following removal of the medium and subsequent cell scraping. Once collected into pre-chilled 2 mL microcentrifuge tubes, cells were centrifuged at 1000 × g for 5 min, and the pellets were stored at –80.0 °C. Pooled QC samples were generated by combining 10 μL of each sample. Metabolite extraction was performed using 1 mL of a single-phase mixture of degassed isopropanol/acetonitrile/water (3:3:2) at −20 °C for 5 min. The tubes were centrifuged for 30 s at 14,000 × g and the supernatant was collected and concentrated to complete dryness. Metabolite profiling was performed using gas chromatography (GC) coupled with time-of-flight mass spectrometry (TOF-MS). A Restek Corporation Rtx-5Sil MS capillary column was used for metabolite separation.

### 2.5: Data Acquisition and Statistical Analyses

The samples were derivatized for GC-TOF-MS analysis (27). Data values are represented as peak heights, comprising the quantification ion (m/z value) with the respective retention index, allowing for increased precision in identifying lower abundant metabolites. Raw reports contained deconvoluted mass spectra, retention indices, unique ions, standard compound identifiers, compound names, class annotations, and KEGG and PubChem compound identifiers (CIDs). SMILES codes were generated using the PubChem Identifier Exchange Service. Univariate analysis involved independent sample t-tests between each pair of groups to test for significantly different metabolites. False discovery rate (FDR) correction for the p-values was used with a p-value < 0.5 to indicate statistical significance. Log_2_FC line plots accounted for fold changes calculated by dividing the mean of the infected groups by that of the non-infected group. Line plots and partial least squares-discriminant analysis (PLS-DA) plots were generated using RStudio. Statistical data, including fold change and p-value, were used to generate network maps using Metamapp (http://metamapp.fiehnlab.ucdavis.edu/ocpu/library/MetaMapp2020/www/#\). An output file was saved as a Cytoscape node attribute file for significantly up-or downregulated metabolites. ChemRICH, which employs Tanimoto substructure chemical similarity coefficients, was used to cluster metabolites into non-overlapping chemical groups. Fold changes for ChemRICH and Metamapp input were calculated by dividing the median of infected groups by the median of non-infected groups (28). Statistically significant chemical classes were determined using ChemRICH by taking the p-values and median fold changes (29). Downregulated and upregulated chemical groups were determined by taking the increased ratio originally on a 0 – +1 scale with 0.5 as the median and converting it to a –1 – +1 scale.

## Results

### HAdV-5 infection resulted in distinct group clustering, while viral genes were expressed across all groups

To confirm viral infection, a subset of the treated samples was evaluated using RT-qPCR. HAdV-5 infection increased early-phase gene expression including E2A and E4 across time points and dosages, confirming viral infection (Figure S1a-b). The evolution of infection groups by PLS-DA revealed that non-infected cells clustered away from the infected cells across all treatments (Figure S2a-d). Notably, one replicate at 12HPI from the 2.0MOI group overlapped with the noninfected cluster, signifying a non-infected outlying sample, which was removed from the downstream analyses. Furthermore, one replicate from the 0.5MOI group at 36HPI failed the injection and was not included in the downstream analyses. Nonetheless, there were subtle differences in the groups of infected cells according to the dose.

### HAdV-5 infection for 36 h disrupts amino acid, lipid, nucleic acid, and sugar metabolism with notable downregulation

After viral infection was confirmed and responses were distinct between the control and treatment groups, we proceeded by classifying the directional changes of metabolites at each time point. A total of 182 individual metabolites were identified in our analysis, and all downstream metabolomic analyses were performed using these specific metabolites. Fold changes between the non-infected cells and the three treatment groups were calculated. Using ChemRICH to evaluate statistical enrichment, multiple classes of metabolism were enriched. The highest enrichment was observed at 6 and 12HPI. Predominant downregulation of metabolic classes was observed at 6HPI and 12HPI (Figure 1). Most amino acids were immediately downregulated across all dosages.

**Figure 1:**
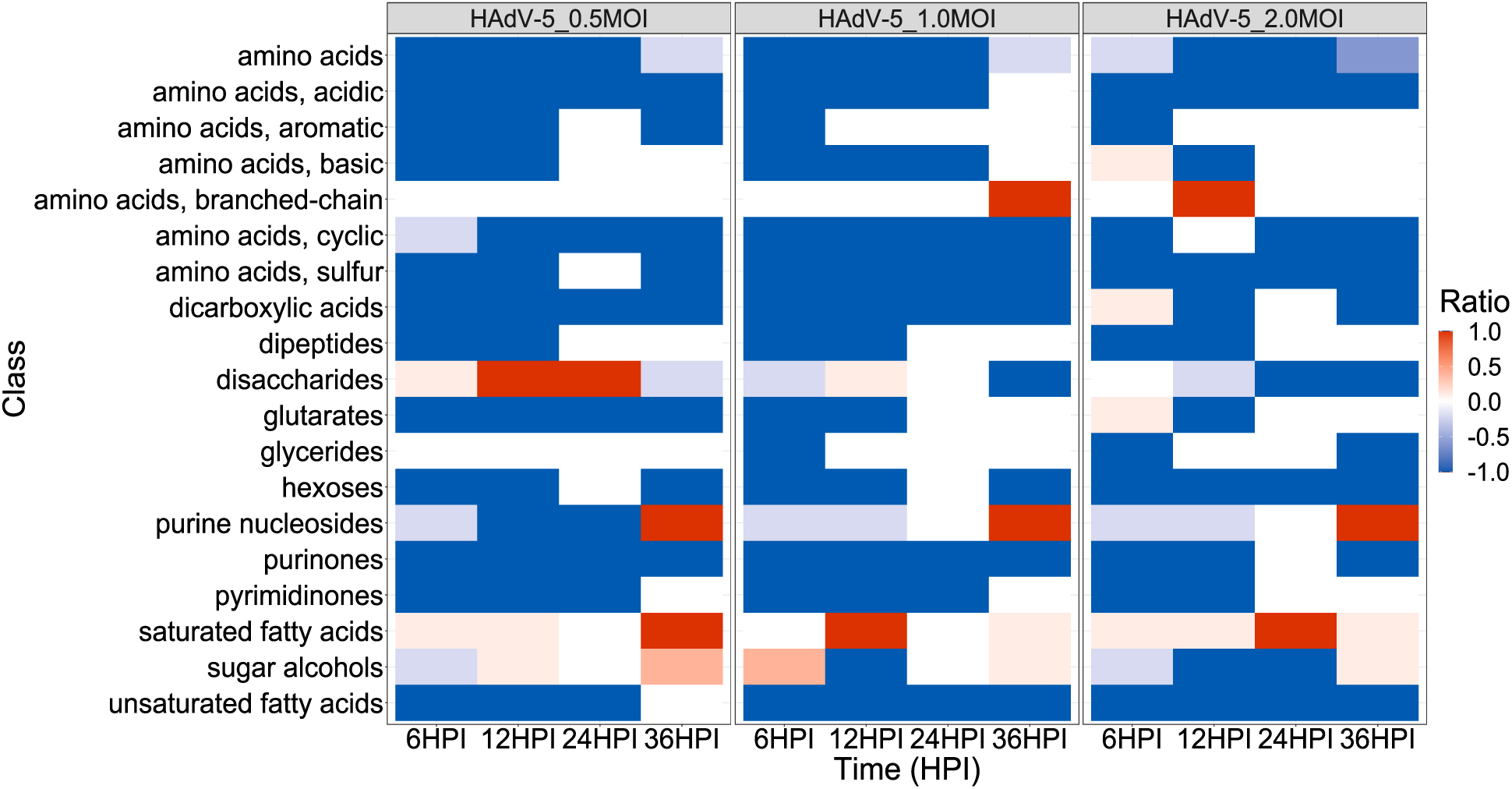
ChemRICH analysis reveals differential metabolite classes between infected and non-infected fibroblasts. Heat map of altered metabolite classes. Metabolite classes are in rows across time and the ratio of up– and downregulation corresponds to the color of each tile. Darker blue indicates greater downregulation, while darker red indicates greater upregulation.

Metabolites involved in unsaturated fatty acid metabolism were also immediately down-regulated. Nucleic acid metabolism also was impacted in early infection, indicated by a downregulation of purine nucleosides and purinones. Notably, additional aspects of metabolism were downregulated, including acidic amino acids, aromatic amino acids, dipeptides, and pyrimidine metabolism (Supplementary Table II-IV). Upregulation was observed in the levels of disaccharides, sugar alcohols, and saturated fatty acid metabolites. Using MetaMapp, we grouped the significantly altered metabolites based on biochemical and mass spectral similarities (34). Across all three dosages, our analyses revealed that HAdV-5 infection disrupted multiple metabolic pathways in the fibroblasts (Figure 1 and S3). The peak metabolic downregulation was observed at 12HPI. The three classes of metabolism related to 6HPI continued to exhibit downregulation, and we continued to see additional significantly altered chemical classes of metabolism (Figure 1, Supplementary Tables V-VII). Across all three dosages, the number of significant metabolites detected at 24 HPI was reduced, suggesting that saturation of the optimal effects of viral infection was present between 12 and 24 HPI. The data at 24HPI revealed fewer altered metabolic categories, with amino acid and unsaturated fatty acid classes shared across all dosages (Figure 1, Supplementary Tables VIII-X). Network analyses revealed consistency in metabolic shifts across metabolite classes (Figure S5). Data at 36HPI revealed upregulation of saturated fatty acids at an MOI of 0.5, and downregulation of unsaturated fatty acid metabolism at 2.0 MOI (Figure 1). Across the three dosages, amino acid and dicarboxylic acid metabolism was downregulated (Supplementary Table XI-XIII). Network analyses of the perturbed metabolites revealed clustering of amino acids, carbohydrates, lipids, and nucleic acid metabolites (Figure S6).

### HAdV-5 infection induces a classical glycolytic interaction that perturbs cysteine, purine, and unsaturated fatty acid metabolism

Adenoviruses typically upregulate glucose metabolism through glycolytic pathways. At our early time points, upregulation of glucose-1-phosphate was observed, and the TCA cycle metabolites fumaric acid and malic acid were downregulated (Figure 2). This coincides with the Warburg effect, whereby metabolic dedication is shifted in favor of glycolysis, as opposed to pathways downstream of the TCA cycle. Additionally, cysteine, purine, and unsaturated fatty acid metabolism were altered. Log_2_FC line plots of cysteine derivatives associated with the *amino acid and sulfur* clusters in the ChemRICH plots revealed prominent downregulation, notably at 6 and 12 HPI, with recovery after 12 HPI (Figure 3). The metabolites included cystathionine, cysteine, homocysteine, and methionine. Similarly, purine derivatives were constrained to purine nucleoside and purinone clusters. Purine metabolites adenine and MTA were upregulated through to 36 HPI, while the purine metabolites, guanine, and hypoxanthine, tended to be downregulated (Figure 4). We also tracked unsaturated fatty acid metabolites and their fold change values over time (Figure 5). The network maps demonstrated that linoleic acid, oleic acid, and palmitoleic acid were downregulated across the time of infection, notably at the 2.0MOI dose, suggesting that long-chain fatty acids may be more susceptible to HAdV-induced alterations in metabolism than medium and short-chain fatty acids.

**Figure 2:**
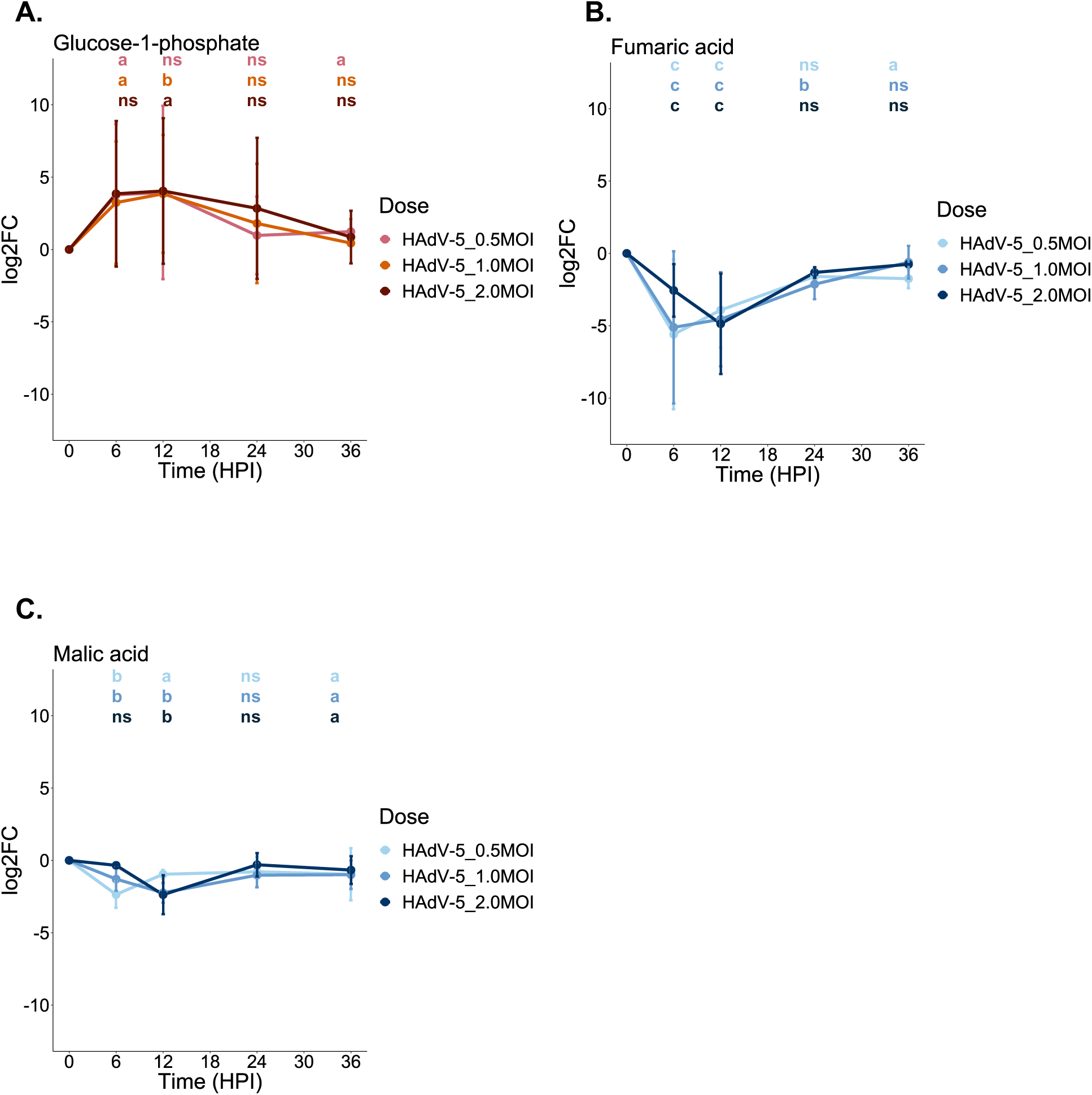
HAdV-5 infection exhibits metabolic shifts consistent with the Warburg effect. **A**. Line plot progression of glucose-1-phosphate log_2_ fold change values relative to non-infected cells. **B.** Line plot progression of fumaric acid log_2_ fold change values relative to non-infected cells. **C.** Line plot progression of malic acid log_2_ fold change values relative to non-infected cells. Error bars indicate the standard error of the mean (SEM). False discovery rate (FDR) correction was used to calculate the p-value. **a** = P < 0.05, **b** = P < 0.01, **c** = P < 0.001, **ns** = not significant.

**Figure 3:**
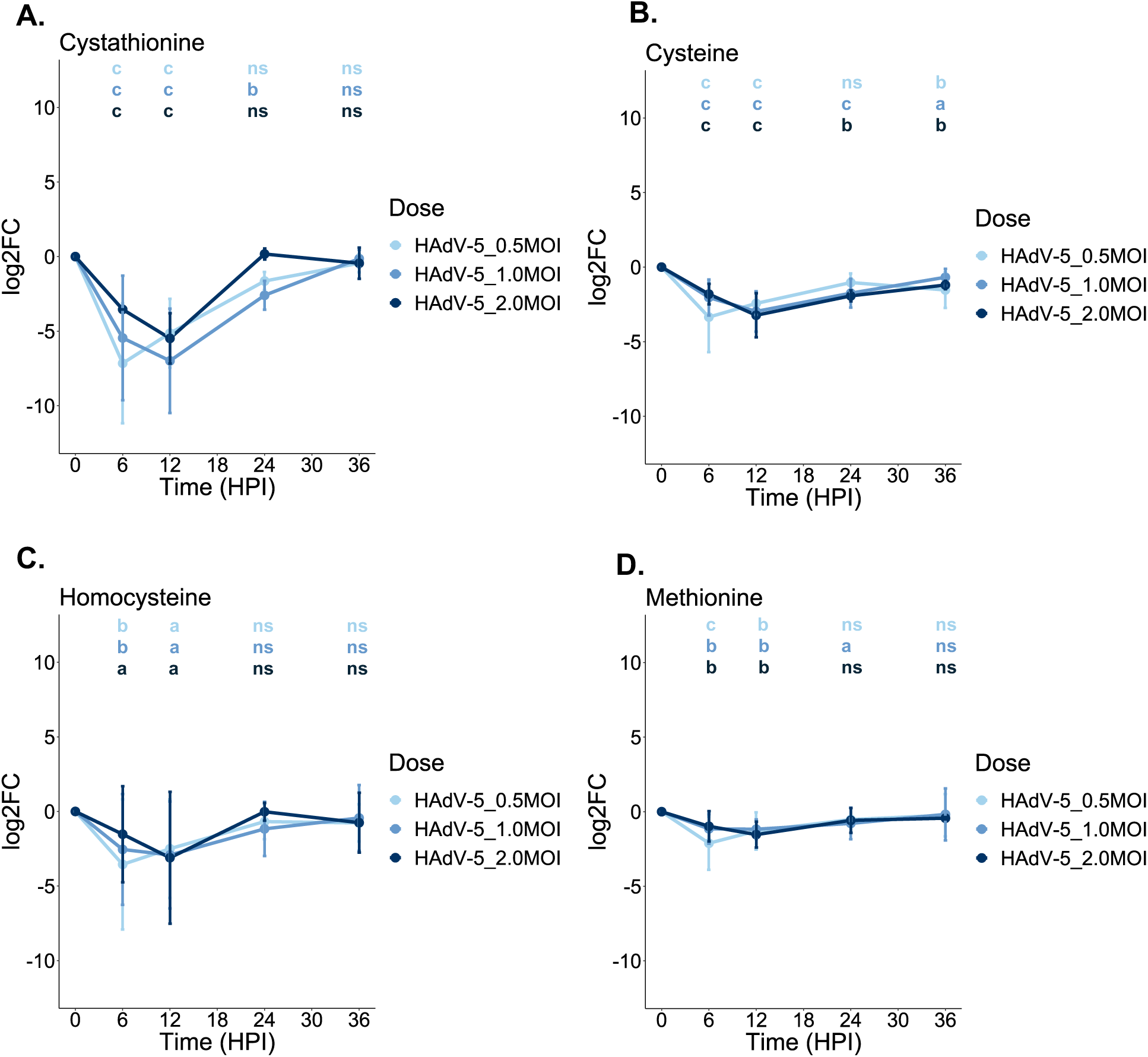
Cysteine metabolites exhibit significant downregulation, notably at 6 and 12 HPI. **A**. Line plot progression of cystathionine log_2_ fold change values relative to non-infected cells. **B.** Line plot progression of cysteine log_2_ fold change values relative to non-infected cells**. C.** Line plot progression of homocysteine log_2_ fold change values relative to non-infected cells. **D.** Line plot progression of methionine log_2_ fold change values relative to non-infected cells. Error bars indicate the standard error of the mean (SEM). False discovery rate (FDR) correction was used to calculate the p-value. **a** = P < 0.05, **b** = P < 0.01, **c** = P < 0.001, **ns** = not significant.

**Figure 4:**
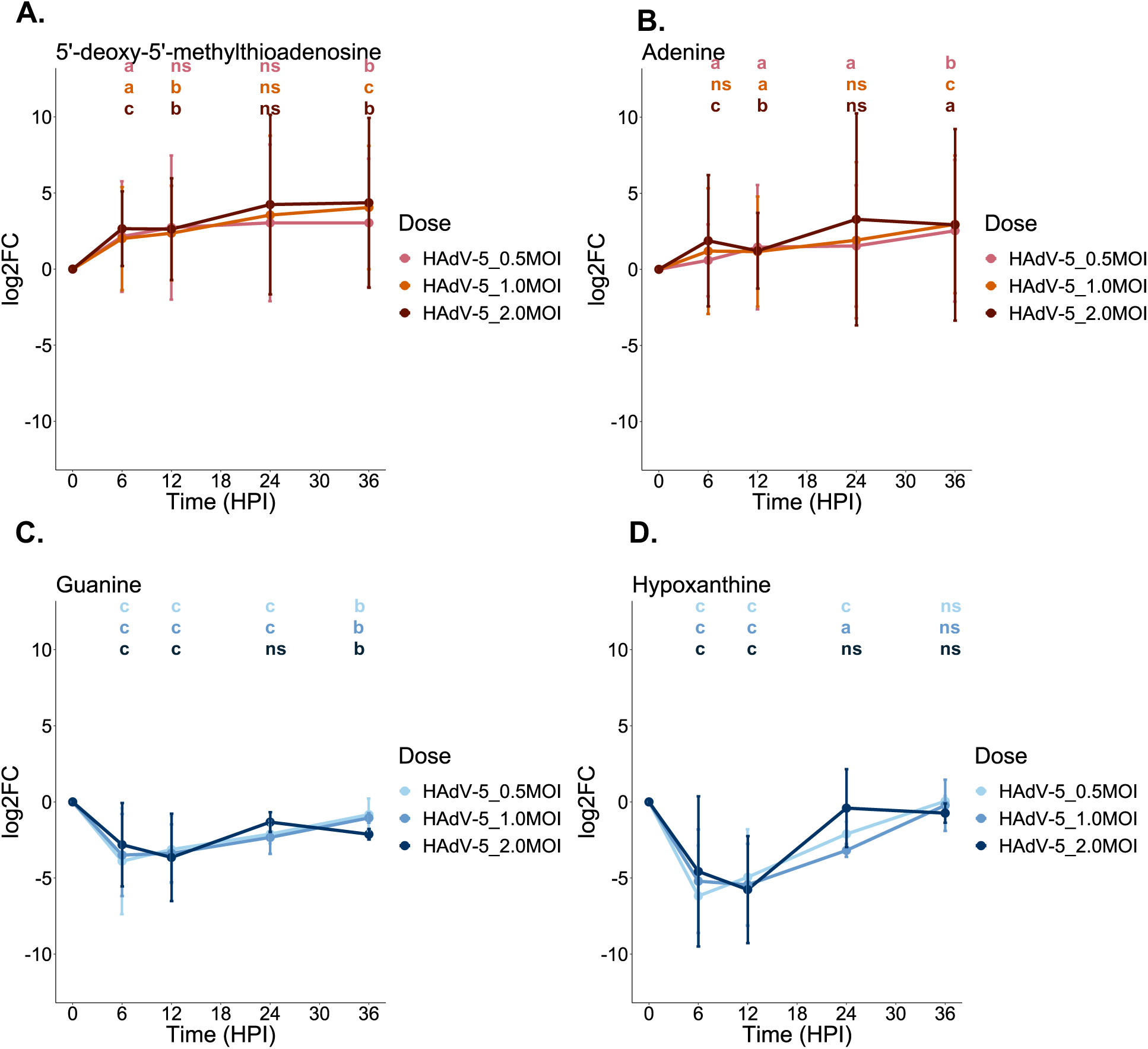
Purine metabolites exhibit dynamic changes over time. **A**. Line plot progression of MTA log_2_ fold change values relative to non-infected cells. **B.** Line plot progression of adenine log_2_ fold change values relative to non-infected cells**. C.** Line plot progression of guanine log_2_ fold change values relative to non-infected cells. **D.** Line plot progression of hypoxanthine log_2_ fold change values relative to non-infected cells. Error bars indicate the standard error of the mean (SEM). False discovery rate (FDR) correction was used to calculate the p-value. **a** = P < 0.05, **b** = P < 0.01, **c** = P < 0.001, **ns** = not significant.

**Figure 5:**
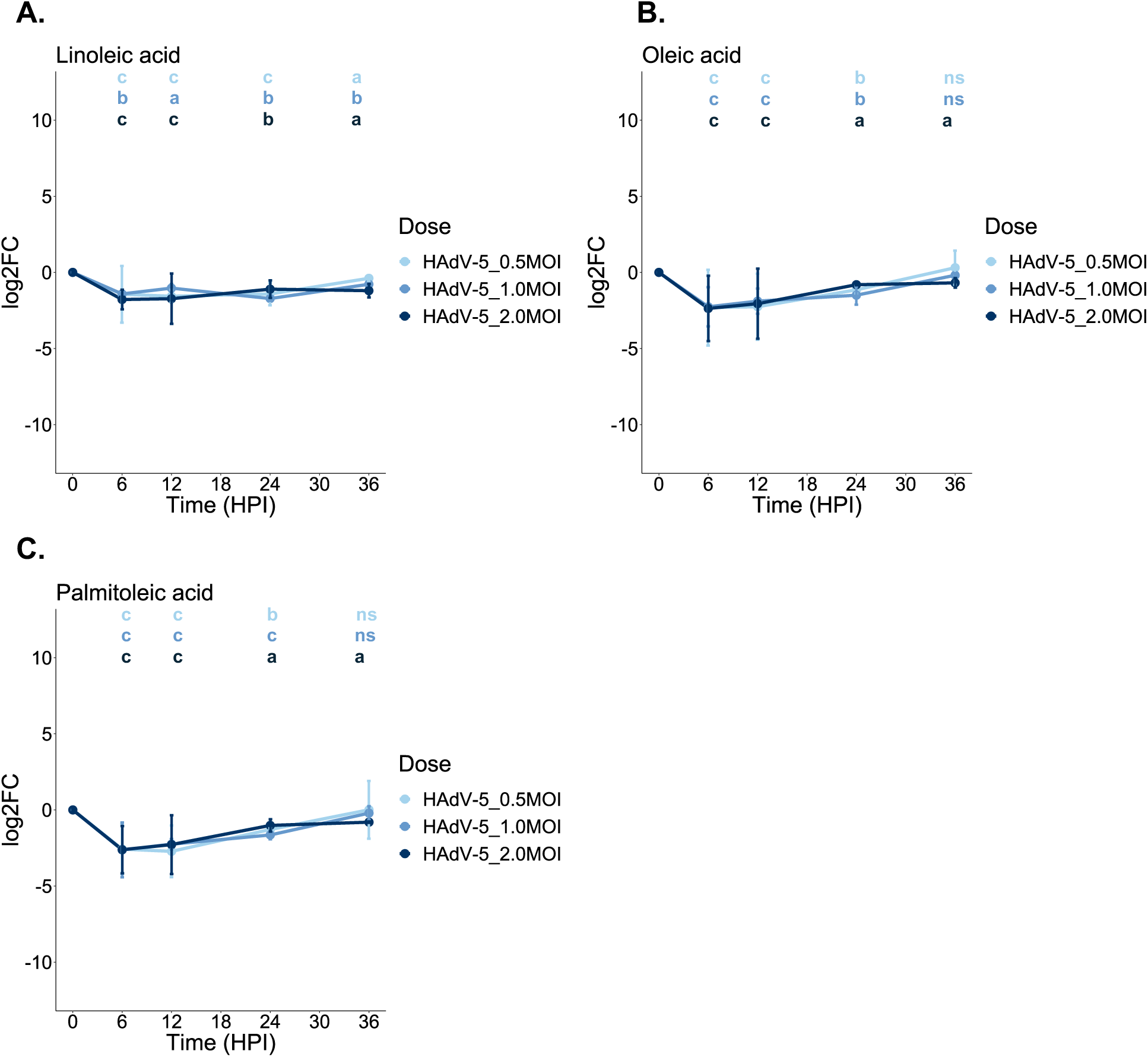
Unsaturated fatty acid metabolites exhibit significant downregulation, notably at 6 and 12 HPI before equilibrating. **A** Line plot progression of linoleic acid log_2_ fold change values relative to non-infected cells. **B** Line plot progression of oleic acid log_2_ fold change values relative to non-infected cells**. C** Line plot progression of palmitoleic acid log_2_ fold change values relative to non-infected cells. Error bars indicate the standard error of the mean (SEM). False discovery rate (FDR) correction was used to calculate the p-value. **a** = P < 0.05, **b** = P < 0.01, **c** = P < 0.001, **ns** = not significant.

## Discussion

The last decade has seen an explosion of studies examining viruses and their global cellular manipulation of host metabolism. The advent of analytical technologies has led to the discovery of new metabolites and their respective pathways that are used by multiple viral species. In the case of human adenoviruses, the mechanisms regulating glycolysis and cellular respiration upon adenoviral infection are now being revealed (35,36). While individual proteins of HAdV-5 have been shown to disrupt cellular metabolism (8,16,17), a more comprehensive examination of the impacted host metabolic pathways in response to an intact adenovirus is lacking. Therefore, we sought to better characterize the intracellular metabolic response to HAdV-5 infection in quiescent fibroblasts, HEK293 cells, using multiple dosages and durations of infection. HEK293 cells have previously been used in studies examining adenoviral disruption of host metabolism, although these studies used more targeted approaches (18,37). In the present study, 182 metabolites were identified, with the majority centered on amino acid, carbohydrate, lipid, and nucleic acid metabolism. We discovered that the greatest metabolic response was as early as 6 HPI across all dosages, with peak metabolic dysregulation observed at 12 HPI, indicating how rapidly a viral infection can disrupt host metabolism independent of the dose of the infection. At 24 HPI, a marked reduction in the number of statistically altered metabolites was observed, which likely reflects the post-peak depletion of metabolic substrates associated with late-phase genetic activation of the adenoviral life cycle. Because most adenoviral studies on the disruption of metabolism utilize a high viral titer at infection or a comparison of high and low doses on a tenfold scale, we purposefully examined the impact of infection on a narrower, more refined (twofold) scale. Interestingly, the profiles of the 0.5, 1.0, and 2.0 MOI-treated groups were similar, suggesting that a minimum threshold to activate a host response can be achieved at a relatively low exposure and in a short period. Nonetheless, unique metabolic responses were observed across dosages and measured time intervals post-infection.

Consistent with previous studies, we observed a typical upregulation of glycolysis, which can be inferred by the upregulation of glycolytic and downregulation of Krebs cycle metabolites. However, the most unique and significant contribution of the present study is the discovery of profound changes in the metabolism of cysteine, purine, and unsaturated fatty acids.

### Cysteine Metabolism

The most notable cysteine metabolites altered throughout the infection timeline were cysteine, cystathionine, homocysteine, hypotaurine, and methionine. In the presence of sufficient methionine, homocysteine is produced, which in turn forms cystathionine through cystathionine β-synthase (CBS) and transsulfuration (38). CBS catalyzes the endogenous synthesis of cystathionine, in which serine undergoes condensation with homocysteine (38,39). Cystathionine is then converted into cysteine by γ-cystathionase. After its formation, cysteine can continue down two pathways. The first involves the formation of glutathione, which is the most prominent, non-enzymatic antioxidant stimulated during redox stress (40–42). Alternatively, cysteine can be converted to cysteine sulfinic acid, which is subsequently decarboxylated to hypotaurine (43–45) (Figure 6).

**Figure 6:**
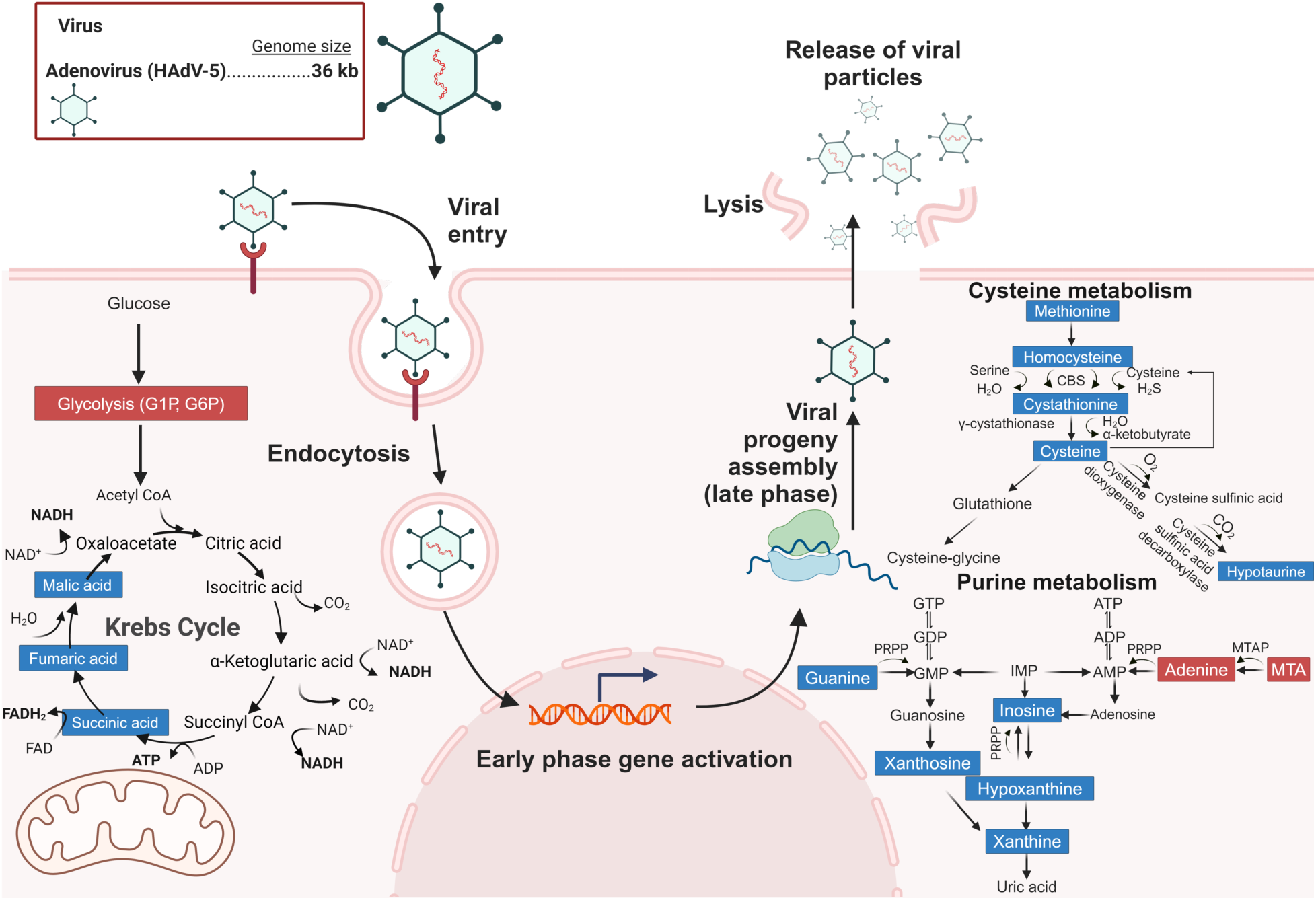
Pathways implicated in this study. Overview of HAdV-5 life cycle with perturbed pathways. The red and blue labels indicate upregulation and downregulation, respectively. Made using BioRender.com. *AMP*, Adenosine monophosphate; *ADP*, Adenosine diphosphate; *ATP*, Adenosine triphosphate; *CBS*, Cystathionine-β-synthase; *FAD*, Flavin adenine dinucleotide; *FADH2*, Flavin adenine dinucleotide (reduced form); *G1P*, Glucose-1-phosphate; *G6P*, Glucose-6-phosphate; *GMP*, Guanosine monophosphate; *GDP*, Guanosine diphosphate; *GTP*, Guanosine triphosphate; *IMP*, Inosine monophosphate; *MTA*, 5’-deoxy-5’methylthioadenosine; *MTAP*, 5’-deoxy-5’methylthioadenosine phosphorylase; *NAD^+^*, Nicotinamide adenine dinucleotide; *NADH*, Nicotinamide adenine dinucleotide (reduced form); *PRPP*, Phosphoribosyl pyrophosphate.

Cystathionine has anti-inflammatory and anti-apoptotic potential in addition to eliminating superoxide radicals (38). In our study, cystathionine was reduced in a dose-dependent manner, suggesting that HAdV infection can compromise a cell’s self-defense against a viral insult by reducing its anti-inflammatory and, ultimately, antioxidant potential. In addition, homocysteine exhibited a trend similar to that of cystathionine. Homocysteine is a sulfhydryl-containing, non-proteinogenic amino acid, essential for cell cycle progression and homeostasis (43). Dysregulation of homocysteine, including hypohomocysteinemia and hyperhomocysteinemia, has been linked to multiple diseases, rendering changes in homocysteine levels as a useful marker for impaired amino acid metabolism depending on the condition (46–48). A meta-analysis of HIV-infected subjects under antiretroviral therapy revealed that mean homocysteine levels were elevated compared to non-treated HIV subjects (49), suggesting that the maintenance of homocysteine levels is important for sustaining cellular health. Homocysteine has also been implicated as a potential biomarker of COVID-19 infection as levels were increased in COVID-19 patients (50,51) and remained elevated for three months in subjects diagnosed with COVID-19-associated pneumonia (50). While our results substantiate the idea that viral infections perturb homocysteine metabolism, resulting in profound changes in levels, they advance our knowledge by indicating that the initial insult may suppress cysteine metabolism. Furthermore, post-infection elevation in homocysteine levels may serve as an indication of cellular rebound to support cell recovery. These results were unique and seminal. Future studies that include targeted examination of enzymes that regulate cysteine metabolism will provide additional insights into whether the changes are due to increased utilization/degradation and/or decreased *de novo* synthesis.

### Purine Metabolism

While serving as structural components of nucleic acids, adenine and guanine may also contribute to the regulation of cellular energetics and multiple signal transduction pathways (52). Cells require purines for their growth, proliferation, and survival. In addition to adenine and guanine, metabolites involved in purine metabolism were significantly altered in our study included 5’-deoxy-5’methylthioadenosine (MTA), inosine, hypoxanthine, and xanthine. Nucleoside inosine is converted to hypoxanthine, which is then converted into xanthine (53–56). Xanthine can also form after the conversion of guanosine into xanthosine. After xanthine synthesis, uric acid results from the oxidation of xanthine (Figure 6) (57), and may possess antioxidant potential. In the present study, adenine and MTA were upregulated across most time points, while guanine, hypoxanthine, and xanthine were downregulated, notably at 6 and 12 HPI. This reciprocal change in adenine and xanthines suggests that ATP utilization is increased and adenine recycling is increased to support the increased energetic burden induced by viral infection. The reduction in xanthines suggests that the antioxidant potential of the host is compromised during adenoviral infection.

### Unsaturated Fatty Acid Metabolism

Unsaturated fatty acids can be divided into two categories: monounsaturated fatty acids (MUFA) and polyunsaturated fatty acids (PUFA). Monounsaturated fatty acids function in apoptosis, cell proliferation and growth, membrane dynamics, and unfolded protein response (58). In the present study, oleic acid was downregulated across all time points. Oleic acid synthesis is achieved when the saturated FA palmitic acid is formed when acetyl CoA and malonyl CoA are converted by fatty acid synthase (59). Palmitic acid is then desaturated to palmitoleic acid or stearic acid, which is another saturated fatty acid. Stearic acid is desaturated to form oleic acid. Palmitoleic acid was downregulated at 6 and 12 HPI. In addition to MUFAs, PUFAs were also downregulated, most notably linoleic acid (18:2) (60). Alpha-linoleic acid’s (aLA) antiviral potential against multiple viral species including Dengue, SARS-CoV-2, and Zika has been reported. Alpha-LA inhibits Zika mRNA production in cells without loss of cell viability, while also interrupting the binding and cellular entry of the virus (61). Therefore, our data on HAdV-induced downregulation of PUFAs suggest that the persistence of a viral infection may be perpetuated by the immediate suppression of key PUFAs during the initial phase. Alternatively, the initial rapid decrease in MUFAs and PUFAs may reflect a rapid shift in increased lipid metabolism to support the increased energetic burden induced by an initial viral insult. Regardless of the purpose, the data suggest that dietary supplements of PUFAs may help alleviate the cellular energy burden and decrease the recovery time.

### Summary

Our results revealed unique metabolic pathways perturbed by adenoviral infection. Given that viral infections are commonly associated with primary disruption of the lung epithelium, our results provide insights into additional sites of cellular metabolism. The data demonstrated the downregulation of TCA cycle intermediates and upregulation of glycolytic intermediates with simultaneous changes in cysteine, purine, and unsaturated fatty acid metabolism. Changes in these metabolic pathways are indicative of an increase in glucose metabolism, likely supporting the rapid increase in cellular energetics induced by a viral insult. The infected cells also showed increased purine (adenine) recycling, which may support the increased energetic burden at the expense of producing uric acid, which may help ameliorate the potential for inflammatory and oxidative injury. The data also suggest that an adenoviral infection induces a shift in cellular priority toward supporting the energetic burden at the expense of increasing the cell’s susceptibility to inflammatory and oxidative injury, at least during the initial and early phases of the infection that persists over the first 24 h. This is supported by the downregulation of unsaturated fatty acid metabolites and a vast number of amino acid metabolites, particularly cysteine pathway intermediates. The doses utilized here demonstrate that adenoviral infections at relatively low doses are sufficient to induce profound effects on host metabolism and help identify a minimal effective dose to reach a metabolic threshold response. The time points used revealed that the response threshold was achieved relatively rapidly, suggesting that more immediate intervention should help ameliorate the duration and impact of the infection and reduce recovery time. We envision that these findings will facilitate the implementation of more mechanistic studies to assist in the development of more robust therapeutic interventions (pharmaceutical, dietary, or otherwise) to combat adenoviral infections.

## Acknowledgments

We thank all members of the Grasis and Ortiz labs for related discussions and collective advice regarding the generation of this manuscript. We are thankful for the metabolomic training and services provided by Dr. Oliver Fiehn and members of the West Coast Metabolomics Center at the University of California, Davis. This research was supported by NSF BII: Host-Virus Evolutionary Dynamics Institute (JAG, 2119968). BJS was supported by a fellowship from a USDA HSI Education Training grant (2021-03397).

---

**Supplemental Table I:**
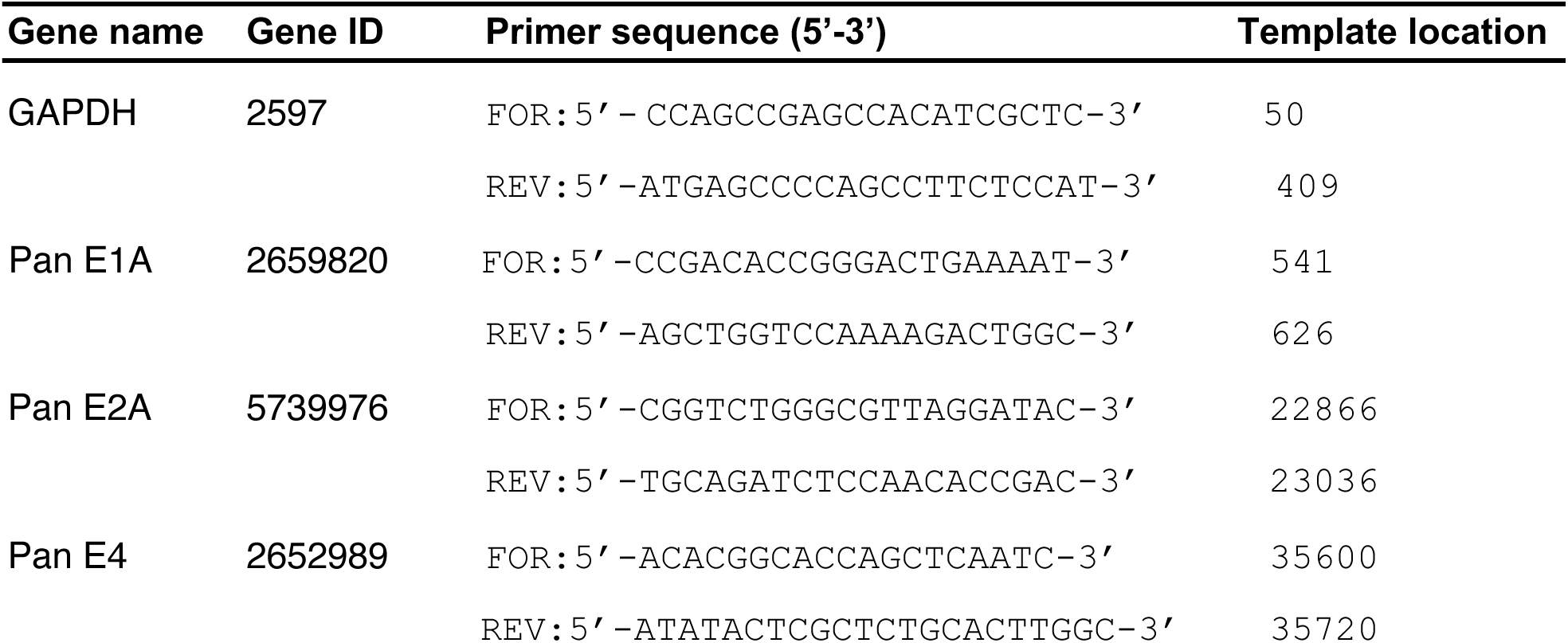
Primers used in RT-qPCR analysis.

**Supplemental Table II:** ChemRICH cluster information for 0.5MOI_6HPI.

| Cluster name | Cluster size | P-value | Transformed P-value | Altered metabolites | Up/down ratio |
| --- | --- | --- | --- | --- | --- |
| adipates | 3 | 5.00E-03 | 9.99E+00 | 2 | 0.1 |
| amino acids | 9 | 4.60E-05 | 2.87E+01 | 6 | -1 |
| amino acids, acidic | 3 | 3.60E-13 | 2.07E+01 | 3 | -1 |
| amino acids, aromatic | 3 | 1.00E-09 | 0.00E+00 | 3 | -1 |
| amino acids, basic | 3 | 1.00E+00 | 0.00E+00 | 1 | -1 |
| amino acids, branched-chain | 3 | 1.00E+00 | 2.85E+01 | 0 | 0 |
| amino acids, cyclic | 4 | 4.40E-13 | 2.21E+01 | 4 | -0.8 |
| amino acids, sulfur | 5 | 2.60E-10 | 0.00E+00 | 5 | -1 |
| benzene derivatives | 3 | 1.00E+00 | 0.00E+00 | 0 | 0 |
| biogenic polyamines | 4 | 1.00E+00 | 0.00E+00 | 1 | -1 |
| cholestenes | 4 | 1.00E+00 | 4.20E+00 | 1 | 1 |
| dicarboxylic acids | 5 | 1.50E-02 | 1.01E+01 | 3 | -1 |
| dipeptides | 5 | 4.30E-05 | 7.90E+00 | 2 | -1 |
| disaccharides | 6 | 3.70E-04 | 0.00E+00 | 4 | 0.1 |
| ethanolamines | 3 | 1.00E+00 | 2.98E+00 | 1 | -1 |
| glutarates | 3 | 5.10E-02 | 0.00E+00 | 2 | -1 |
| glyceric acids | 3 | 1.00E+00 | 0.00E+00 | 0 | 0 |
| glycerides | 3 | 1.00E+00 | 1.21E+01 | 0 | 0 |
| hexosephosphates | 6 | 5.80E-06 | 0.00E+00 | 2 | 0.1 |
| hexoses | 3 | 1.00E+00 | 3.12E+00 | 1 | -1 |
| pentoses | 3 | 4.40E-02 | 7.31E+00 | 2 | -1 |
| purine nucleosides | 7 | 6.70E-04 | 1.53E+01 | 5 | -0.8 |
| purinones | 3 | 2.20E-07 | 3.24E+00 | 3 | -1 |
| pyrimidine nucleosides | 3 | 3.90E-02 | 3.30E+00 | 2 | -1 |
| pyrimidinones | 3 | 3.70E-02 | 7.26E+00 | 2 | -1 |
| saturated fatty acids | 14 | 7.00E-04 | 2.96E+00 | 2 | 0.1 |
| sugar alcohols | 9 | 5.20E-02 | 9.85E+00 | 3 | -0.6 |
| unsaturated fatty acids | 6 | 5.30E-05 | 9.99E+00 | 5 | -1 |

**Supplemental Table III:** ChemRICH cluster information for 1.0MOI_6HPI.

| Cluster name | Cluster size | P-value | Transformed P-value | Altered metabolites | Up/down ratio |
| --- | --- | --- | --- | --- | --- |
| adipates | 3 | 1.00E+00 | 0.00E+00 | 1 | -1 |
| amino acids | 8 | 1.30E-02 | 4.34E+00 | 3 | -1 |
| amino acids, acidic | 3 | 3.10E-14 | 3.11E+01 | 3 | -1 |
| amino acids, aromatic | 3 | 1.00E-06 | 1.38E+01 | 3 | -1 |
| amino acids, basic | 3 | 1.00E+00 | 0.00E+00 | 1 | -1 |
| amino acids, branched-chain | 3 | 1.00E+00 | 0.00E+00 | 0 | 0 |
| amino acids, cyclic | 4 | 4.00E-03 | 5.52E+00 | 3 | -1 |
| amino acids, sulfur | 4 | 2.60E-10 | 2.21E+01 | 4 | -1 |
| benzene derivatives | 3 | 1.00E+00 | 0.00E+00 | 1 | 1 |
| biogenic polyamines | 4 | 1.00E+00 | 0.00E+00 | 1 | -1 |
| cholestenes | 4 | 8.20E-03 | 4.80E+00 | 3 | 1 |
| dicarboxylic acids | 5 | 1.60E-01 | 1.83E+00 | 2 | -1 |
| dipeptides | 5 | 1.00E+00 | 0.00E+00 | 1 | -1 |
| disaccharides | 6 | 9.50E-03 | 4.66E+00 | 3 | -0.6 |
| ethanolamines | 3 | 1.00E+00 | 0.00E+00 | 1 | -1 |
| glutarates | 3 | 1.00E+00 | 0.00E+00 | 1 | -1 |
| glyceric acids | 3 | 1.00E+00 | 0.00E+00 | 0 | 0 |
| glycerides | 3 | 1.00E+00 | 0.00E+00 | 1 | -1 |
| hexosephosphates | 6 | 8.20E-06 | 1.17E+01 | 3 | 0.4 |
| hexoses | 3 | 1.00E+00 | 0.00E+00 | 1 | -1 |
| pentoses | 3 | 1.00E+00 | 0.00E+00 | 0 | 0 |
| purine nucleosides | 7 | 1.30E-02 | 4.34E+00 | 4 | -0.8 |
| purinones | 3 | 2.20E-07 | 1.53E+01 | 3 | -1 |
| pyrimidine nucleosides | 3 | 3.90E-02 | 3.24E+00 | 2 | -1 |
| pyrimidinones | 3 | 3.70E-02 | 3.30E+00 | 2 | -1 |
| saturated fatty acids | 14 | 1.00E+00 | 0.00E+00 | 0 | 0 |
| sugar alcohols | 9 | 1.40E-02 | 4.27E+00 | 3 | 0.4 |
| unsaturated fatty acids | 6 | 2.00E-03 | 6.21E+00 | 4 | -1 |

**Supplemental Table IV:** ChemRICH cluster information for 2.0MOI_6HPI.

| Cluster name | Cluster size | P-value | Transformed P-value | Altered metabolites | Up/down ratio |
| --- | --- | --- | --- | --- | --- |
| adipates | 3 | 1.00E+00 | 0.00E+00 | 1 | -1 |
| amino acids | 9 | 2.70E-02 | 3.61E+00 | 4 | -0.6 |
| amino acids, acidic | 3 | 4.40E-11 | 2.38E+01 | 3 | -1 |
| amino acids, aromatic | 3 | 4.10E-02 | 3.19E+00 | 2 | -1 |
| amino acids, basic | 3 | 6.40E-04 | 7.35E+00 | 2 | 0.1 |
| amino acids, branched-chain | 3 | 1.00E+00 | 0.00E+00 | 0 | 0 |
| amino acids, cyclic | 4 | 4.00E-03 | 5.52E+00 | 3 | -1 |
| amino acids, sulfur | 5 | 9.80E-06 | 1.15E+01 | 4 | -1 |
| benzene derivatives | 3 | 1.00E+00 | 0.00E+00 | 0 | 0 |
| biogenic polyamines | 4 | 1.00E+00 | 0.00E+00 | 1 | -1 |
| cholestenes | 4 | 6.10E-03 | 5.10E+00 | 3 | 1 |
| dicarboxylic acids | 5 | 9.50E-02 | 2.35E+00 | 2 | 0.1 |
| dipeptides | 5 | 1.00E+00 | 0.00E+00 | 1 | -1 |
| disaccharides | 6 | 1.00E+00 | 0.00E+00 | 0 | 0 |
| ethanolamines | 3 | 1.00E+00 | 0.00E+00 | 1 | -1 |
| glutarates | 3 | 5.20E-02 | 2.96E+00 | 2 | 0.1 |
| glyceric acids | 3 | 1.00E+00 | 0.00E+00 | 0 | 0 |
| glycerides | 3 | 1.00E+00 | 0.00E+00 | 1 | -1 |
| hexosephosphates | 6 | 1.10E-03 | 6.81E+00 | 2 | 0.1 |
| hexoses | 3 | 3.90E-02 | 3.24E+00 | 2 | -1 |
| pentoses | 3 | 1.00E+00 | 0.00E+00 | 0 | 0 |
| purine nucleosides | 7 | 1.60E-05 | 1.10E+01 | 5 | -0.8 |
| purinones | 3 | 2.20E-07 | 1.53E+01 | 3 | -1 |
| pyrimidine nucleosides | 3 | 3.90E-02 | 3.24E+00 | 2 | -1 |
| pyrimidinones | 3 | 1.00E+00 | 0.00E+00 | 1 | -1 |
| saturated fatty acids | 14 | 1.20E-02 | 4.42E+00 | 2 | 0.1 |
| sugar alcohols | 9 | 6.80E-03 | 4.99E+00 | 3 | -0.6 |
| unsaturated fatty acids | 6 | 5.30E-05 | 9.85E+00 | 5 | -1 |

**Supplemental Table V:** ChemRICH cluster information for 0.5MOI_12HPI.

| Cluster name | Cluster size | P-value | Transformed P-value | Altered metabolites | Up/down ratio |
| --- | --- | --- | --- | --- | --- |
| adipates | 3 | 8.00E-04 | 4.34E+00 | 5 | 0.1 |
| amino acids | 9 | 1.30E-02 | 2.07E+01 | 3 | -1 |
| amino acids, acidic | 3 | 1.00E-09 | 0.00E+00 | 1 | -1 |
| amino acids, aromatic | 3 | 1.00E+00 | 0.00E+00 | 1 | -1 |
| amino acids, basic | 3 | 1.00E+00 | 0.00E+00 | 0 | -1 |
| amino acids, branched-chain | 3 | 1.00E+00 | 6.12E+00 | 3 | 0 |
| amino acids, cyclic | 4 | 2.20E-03 | 1.77E+01 | 5 | -1 |
| amino acids, sulfur | 5 | 2.10E-08 | 0.00E+00 | 0 | -1 |
| benzene derivatives | 3 | 1.00E+00 | 2.21E+00 | 2 | 0 |
| biogenic polyamines | 4 | 1.10E-01 | 0.00E+00 | 0 | 0.1 |
| cholestenes | 4 | 1.00E+00 | 0.00E+00 | 1 | 0 |
| dicarboxylic acids | 5 | 1.00E+00 | 4.92E+00 | 2 | -1 |
| dipeptides | 5 | 7.30E-03 | 0.00E+00 | 1 | -1 |
| disaccharides | 6 | 1.00E+00 | 0.00E+00 | 1 | 1 |
| ethanolamines | 3 | 1.00E+00 | 0.00E+00 | 1 | -1 |
| glutarates | 3 | 1.00E+00 | 0.00E+00 | 0 | -1 |
| glyceric acids | 3 | 1.00E+00 | 0.00E+00 | 0 | 0 |
| glycerides | 3 | 1.00E+00 | 0.00E+00 | 1 | 0 |
| hexosephosphates | 6 | 1.00E+00 | 0.00E+00 | 1 | -1 |
| hexoses | 3 | 1.00E+00 | 0.00E+00 | 1 | -1 |
| pentoses | 3 | 1.00E+00 | 2.78E+00 | 3 | -1 |
| purine nucleosides | 7 | 6.20E-02 | 1.33E+01 | 3 | -1 |
| purinones | 3 | 1.70E-06 | 3.22E+00 | 2 | -1 |
| pyrimidine nucleosides | 3 | 4.00E-02 | 2.94E+00 | 2 | -1 |
| pyrimidinones | 3 | 5.30E-02 | 5.20E+00 | 4 | -1 |
| saturated fatty acids | 14 | 5.50E-03 | 9.94E-01 | 2 | 0.1 |
| sugar alcohols | 9 | 3.70E-01 | 1.04E+01 | 5 | 0.1 |
| unsaturated fatty acids | 6 | 3.10E-05 | 4.34E+00 | 5 | -1 |

**Supplemental Table VI:** ChemRICH cluster information for 1.0MOI_12HPI.

| Cluster name | Cluster size | P-value | Transformed P-value | Altered metabolites | Up/down ratio |
| --- | --- | --- | --- | --- | --- |
| adipates | 3 | 2.90E-03 | 5.84E+00 | 5 | -1 |
| amino acids | 9 | 7.80E-16 | 3.48E+01 | 3 | -1 |
| amino acids, acidic | 3 | 1.00E+00 | 0.00E+00 | 0 | 0 |
| amino acids, aromatic | 3 | 1.00E+00 | 0.00E+00 | 1 | -1 |
| amino acids, basic | 3 | 1.00E+00 | 0.00E+00 | 0 | 0 |
| amino acids, branched-chain | 3 | 2.20E-03 | 6.12E+00 | 3 | -1 |
| amino acids, cyclic | 4 | 6.60E-09 | 1.88E+01 | 5 | -1 |
| amino acids, sulfur | 5 | 1.00E+00 | 0.00E+00 | 0 | 0 |
| benzene derivatives | 3 | 1.00E+00 | 0.00E+00 | 1 | -1 |
| biogenic polyamines | 4 | 1.00E+00 | 0.00E+00 | 1 | 1 |
| cholestenes | 4 | 7.00E-02 | 2.66E+00 | 2 | -1 |
| dicarboxylic acids | 5 | 5.60E-05 | 9.79E+00 | 4 | -1 |
| dipeptides | 5 | 6.80E-04 | 7.29E+00 | 2 | 0.1 |
| disaccharides | 6 | 1.00E+00 | 0.00E+00 | 1 | -1 |
| ethanolamines | 3 | 3.70E-02 | 3.30E+00 | 2 | -1 |
| glutarates | 3 | 1.00E+00 | 0.00E+00 | 0 | 0 |
| glyceric acids | 3 | 1.00E+00 | 0.00E+00 | 0 | 0 |
| glycerides | 3 | 1.00E-02 | 4.61E+00 | 3 | 0.4 |
| hexosephosphates | 6 | 1.00E+00 | 0.00E+00 | 1 | -1 |
| hexoses | 3 | 1.00E+00 | 0.00E+00 | 1 | -1 |
| pentoses | 3 | 5.80E-04 | 7.45E+00 | 5 | -0.6 |
| purine nucleosides | 7 | 1.70E-06 | 1.33E+01 | 3 | -1 |
| purinones | 3 | 4.40E-04 | 7.73E+00 | 2 | -1 |
| pyrimidine nucleosides | 3 | 3.80E-02 | 3.27E+00 | 2 | -1 |
| pyrimidinones | 3 | 4.20E-02 | 3.17E+00 | 2 | 1 |
| saturated fatty acids | 14 | 1.00E+00 | 0.00E+00 | 1 | -1 |
| sugar alcohols | 9 | 2.20E-04 | 8.42E+00 | 4 | -1 |
| unsaturated fatty acids | 6 | 2.90E-03 | 5.84E+00 | 5 | -1 |

**Supplemental Table VII:** ChemRICH cluster information for 2.0MOI_12HPI.

| Cluster name | Cluster size | P-value | Transformed P-value | Altered metabolites | Up/down ratio |
| --- | --- | --- | --- | --- | --- |
| adipates | 3 | 1.00E+00 | 0.00E+00 | 1 | -1 |
| amino acids | 9 | 3.40E-03 | 5.68E+00 | 5 | -1 |
| amino acids, acidic | 3 | 6.60E-15 | 3.27E+01 | 3 | -1 |
| amino acids, aromatic | 3 | 1.00E+00 | 0.00E+00 | 0 | 0 |
| amino acids, basic | 3 | 1.00E+00 | 0.00E+00 | 1 | -1 |
| amino acids, branched-chain | 3 | 1.00E+00 | 0.00E+00 | 1 | 1 |
| amino acids, cyclic | 4 | 1.50E-04 | 8.80E+00 | 3 | -1 |
| amino acids, sulfur | 4 | 1.00E-08 | 1.84E+01 | 4 | -1 |
| benzene derivatives | 3 | 1.00E+00 | 0.00E+00 | 0 | 0 |
| biogenic polyamines | 4 | 1.00E-01 | 2.30E+00 | 2 | 0.1 |
| cholestenes | 4 | 1.00E+00 | 0.00E+00 | 1 | 1 |
| dicarboxylic acids | 5 | 8.40E-03 | 4.78E+00 | 3 | -1 |
| dipeptides | 5 | 1.50E-05 | 1.11E+01 | 2 | -1 |
| disaccharides | 6 | 8.60E-03 | 4.76E+00 | 3 | -0.4 |
| ethanolamines | 3 | 2.50E-04 | 8.29E+00 | 2 | -1 |
| glutarates | 3 | 2.20E-02 | 3.82E+00 | 2 | -1 |
| glyceric acids | 3 | 1.00E+00 | 0.00E+00 | 0 | 0 |
| glycerides | 3 | 1.00E+00 | 0.00E+00 | 0 | 0 |
| hexosephosphates | 6 | 2.50E-03 | 5.99E+00 | 2 | 0.1 |
| hexoses | 3 | 1.00E+00 | 0.00E+00 | 1 | -1 |
| pentoses | 3 | 1.00E+00 | 0.00E+00 | 1 | -1 |
| purine nucleosides | 7 | 5.70E-04 | 7.47E+00 | 5 | -0.6 |
| purinones | 3 | 2.70E-06 | 1.28E+01 | 3 | -1 |
| pyrimidine nucleosides | 3 | 3.90E-02 | 3.24E+00 | 2 | -1 |
| pyrimidinones | 3 | 3.90E-02 | 3.24E+00 | 2 | -1 |
| saturated fatty acids | 14 | 1.10E-02 | 4.51E+00 | 2 | 0.1 |
| sugar alcohols | 9 | 1.00E+00 | 0.00E+00 | 1 | -1 |
| unsaturated fatty acids | 6 | 3.80E-05 | 1.02E+01 | 5 | -1 |

**Supplemental Table VIII:** ChemRICH cluster information for 0.5MOI_24HPI.

| Cluster name | Cluster size | P-value | Transformed P-value | Altered metabolites | Up/down ratio |
| --- | --- | --- | --- | --- | --- |
| amino acids | 8 | 1.50E-01 | 1.90E+00 | 2 | -1 |
| amino acids, acidic | 3 | 1.00E+00 | 0.00E+00 | 1 | -1 |
| amino acids, aromatic | 3 | 1.00E+00 | 0.00E+00 | 0 | 0 |
| amino acids, basic | 3 | 1.00E+00 | 0.00E+00 | 0 | 0 |
| amino acids, branched-chain | 3 | 1.00E+00 | 0.00E+00 | 0 | 0 |
| amino acids, cyclic | 3 | 1.00E+00 | 0.00E+00 | 1 | -1 |
| amino acids, sulfur | 5 | 1.00E+00 | 0.00E+00 | 0 | 0 |
| dicarboxylic acids | 7 | 1.00E+00 | 0.00E+00 | 1 | -1 |
| dipeptides | 3 | 1.00E+00 | 0.00E+00 | 0 | 0 |
| disaccharides | 5 | 1.00E+00 | 0.00E+00 | 1 | 1 |
| glutarates | 3 | 1.00E+00 | 0.00E+00 | 1 | -1 |
| glycerides | 3 | 1.00E+00 | 0.00E+00 | 0 | 0 |
| hexoses | 3 | 1.00E+00 | 0.00E+00 | 0 | 0 |
| purine nucleosides | 6 | 1.00E+00 | 0.00E+00 | 1 | -1 |
| purinones | 3 | 6.40E-05 | 9.66E+00 | 3 | -1 |
| pyrimidinones | 3 | 1.00E+00 | 0.00E+00 | 1 | -1 |
| saturated fatty acids | 9 | 1.00E+00 | 0.00E+00 | 0 | 0 |
| sugar acids | 3 | 1.00E+00 | 0.00E+00 | 0 | 0 |
| sugar alcohols | 7 | 1.00E+00 | 0.00E+00 | 0 | 0 |
| unsaturated fatty acids | 4 | 1.90E-03 | 6.27E+00 | 3 | -1 |

**Supplemental Table IX:**
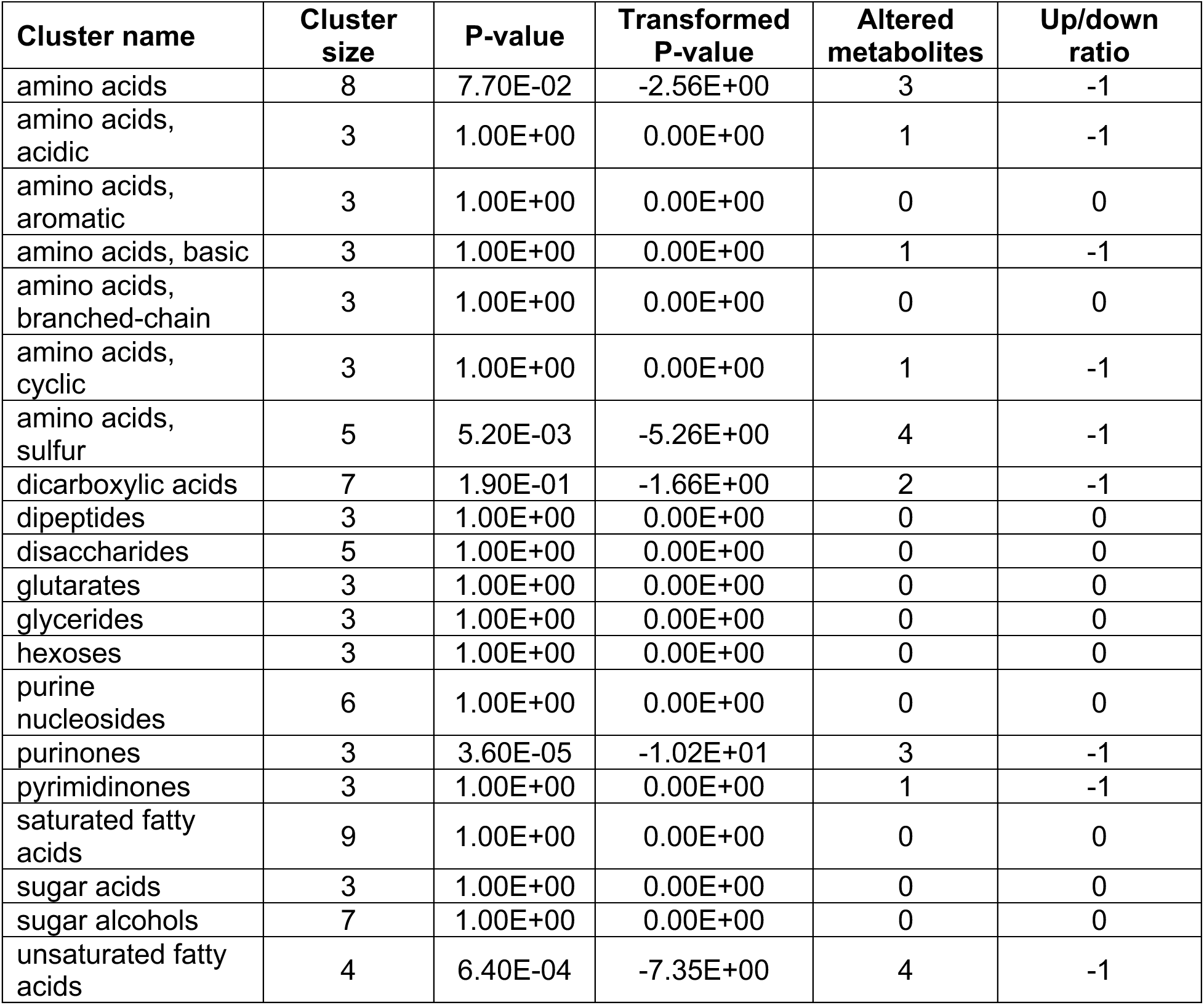
ChemRICH cluster information for 1.0MOI_24HPI.

| Cluster name | Cluster size | P-value | Transformed P-value | Altered metabolites | Up/down ratio |
| --- | --- | --- | --- | --- | --- |
| amino acids | 8 | 7.70E-02 | -2.56E+00 | 3 | -1 |
| amino acids, acidic | 3 | 1.00E+00 | 0.00E+00 | 1 | -1 |
| amino acids, aromatic | 3 | 1.00E+00 | 0.00E+00 | 0 | 0 |
| amino acids, basic | 3 | 1.00E+00 | 0.00E+00 | 1 | -1 |
| amino acids, branched-chain | 3 | 1.00E+00 | 0.00E+00 | 0 | 0 |
| amino acids, cyclic | 3 | 1.00E+00 | 0.00E+00 | 1 | -1 |
| amino acids, sulfur | 5 | 5.20E-03 | -5.26E+00 | 4 | -1 |
| dicarboxylic acids | 7 | 1.90E-01 | -1.66E+00 | 2 | -1 |
| dipeptides | 3 | 1.00E+00 | 0.00E+00 | 0 | 0 |
| disaccharides | 5 | 1.00E+00 | 0.00E+00 | 0 | 0 |
| glutarates | 3 | 1.00E+00 | 0.00E+00 | 0 | 0 |
| glycerides | 3 | 1.00E+00 | 0.00E+00 | 0 | 0 |
| hexoses | 3 | 1.00E+00 | 0.00E+00 | 0 | 0 |
| purine nucleosides | 6 | 1.00E+00 | 0.00E+00 | 0 | 0 |
| purinones | 3 | 3.60E-05 | -1.02E+01 | 3 | -1 |
| pyrimidinones | 3 | 1.00E+00 | 0.00E+00 | 1 | -1 |
| saturated fatty acids | 9 | 1.00E+00 | 0.00E+00 | 0 | 0 |
| sugar acids | 3 | 1.00E+00 | 0.00E+00 | 0 | 0 |
| sugar alcohols | 7 | 1.00E+00 | 0.00E+00 | 0 | 0 |
| unsaturated fatty acids | 4 | 6.40E-04 | -7.35E+00 | 4 | -1 |

**Supplemental Table X:** ChemRICH cluster information for 2.0MOI_24HPI.

| Cluster name | Cluster size | P-value | Transformed P-value | Altered metabolites | Up/down ratio |
| --- | --- | --- | --- | --- | --- |
| amino acids | 8 | 7.70E-02 | 2.85E+00 | 2 | -1 |
| amino acids, acidic | 3 | 1.00E+00 | 3.02E+00 | 2 | -1 |
| amino acids, aromatic | 3 | 1.00E+00 | 0.00E+00 | 0 | 0 |
| amino acids, basic | 3 | 1.00E+00 | 0.00E+00 | 0 | 0 |
| amino acids, branched-chain | 3 | 1.00E+00 | 0.00E+00 | 0 | 0 |
| amino acids, cyclic | 3 | 1.00E+00 | 0.00E+00 | 1 | -1 |
| amino acids, sulfur | 5 | 5.20E-03 | 0.00E+00 | 1 | -1 |
| dicarboxylic acids | 7 | 1.90E-01 | 0.00E+00 | 0 | 0 |
| dipeptides | 3 | 1.00E+00 | 0.00E+00 | 0 | 0 |
| disaccharides | 5 | 1.00E+00 | 0.00E+00 | 1 | -1 |
| glutarates | 3 | 1.00E+00 | 0.00E+00 | 0 | 0 |
| glycerides | 3 | 1.00E+00 | 0.00E+00 | 0 | 0 |
| hexoses | 3 | 1.00E+00 | 0.00E+00 | 1 | -1 |
| purine nucleosides | 6 | 1.00E+00 | 0.00E+00 | 0 | 0 |
| purinones | 3 | 3.60E-05 | 0.00E+00 | 0 | 0 |
| pyrimidinones | 3 | 1.00E+00 | 0.00E+00 | 0 | 0 |
| saturated fatty acids | 9 | 1.00E+00 | 0.00E+00 | 1 | 1 |
| sugar acids | 3 | 1.00E+00 | 0.00E+00 | 0 | 0 |
| sugar alcohols | 7 | 1.00E+00 | 0.00E+00 | 1 | -1 |
| unsaturated fatty acids | 4 | 6.40E-04 | 5.08E+00 | 3 | -1 |

**Supplemental Table XI:** ChemRICH cluster information for 0.5MOI_36HPI.

| Cluster name | Cluster size | P-value | Transformed P-value | Altered metabolites | Up/down ratio |
| --- | --- | --- | --- | --- | --- |
| adipates | 3 | 1.00E+00 | 0.00E+00 | 1 | -1 |
| amino acids | 9 | 4.10E-05 | 1.01E+01 | 6 | -0.4 |
| amino acids, acidic | 3 | 1.00E+00 | 0.00E+00 | 1 | -1 |
| amino acids, aromatic | 3 | 1.00E+00 | 0.00E+00 | 1 | -1 |
| amino acids, basic | 3 | 1.00E+00 | 0.00E+00 | 0 | 0 |
| amino acids, branched-chain | 3 | 1.00E+00 | 0.00E+00 | 0 | 0 |
| amino acids, cyclic | 4 | 1.00E+00 | 0.00E+00 | 1 | -1 |
| amino acids, sulfur | 5 | 1.00E+00 | 0.00E+00 | 2 | -1 |
| benzene derivatives | 3 | 1.00E+00 | 0.00E+00 | 0 | 0 |
| biogenic polyamines | 4 | 1.00E+00 | 0.00E+00 | 1 | -1 |
| cholestenes | 4 | 1.00E+00 | 0.00E+00 | 1 | 1 |
| dicarboxylic acids | 5 | 7.30E-03 | 4.92E+00 | 3 | -1 |
| dipeptides | 5 | 1.00E+00 | 0.00E+00 | 0 | 0 |
| disaccharides | 6 | 6.00E-02 | 2.81E+00 | 3 | -0.4 |
| ethanolamines | 3 | 1.00E+00 | 0.00E+00 | 0 | 0 |
| glutarates | 3 | 1.00E+00 | 0.00E+00 | 1 | -1 |
| glyceric acids | 3 | 1.00E+00 | 0.00E+00 | 0 | 0 |
| glycerides | 3 | 1.00E+00 | 0.00E+00 | 0 | 0 |
| hexosephosphates | 6 | 1.00E+00 | 0.00E+00 | 1 | 1 |
| hexoses | 3 | 1.00E+00 | 0.00E+00 | 1 | -1 |
| pentoses | 3 | 1.00E+00 | 0.00E+00 | 0 | 0 |
| purine nucleosides | 7 | 1.00E+00 | 0.00E+00 | 1 | 1 |
| purinones | 3 | 1.00E+00 | 0.00E+00 | 1 | -1 |
| pyrimidine nucleosides | 3 | 1.00E+00 | 0.00E+00 | 0 | 0 |
| pyrimidinones | 3 | 1.00E+00 | 0.00E+00 | 0 | 0 |
| saturated fatty acids | 14 | 3.10E-02 | 3.47E+00 | 4 | 1 |
| sugar alcohols | 9 | 4.50E-02 | 3.10E+00 | 3 | 0.4 |
| unsaturated fatty acids | 6 | 1.00E+00 | 0.00E+00 | 0 | 0 |

**Supplemental Table XII:** ChemRICH cluster information for 1.0MOI_36HPI.

| Cluster name | Cluster size | P-value | Transformed P-value | Altered metabolites | Up/down ratio |
| --- | --- | --- | --- | --- | --- |
| adipates | 3 | 1.00E+00 | 0.00E+00 | 4 | -1 |
| amino acids | 9 | 1.20E-02 | 4.42E+00 | 0 | -0.6 |
| amino acids, acidic | 3 | 1.00E+00 | 0.00E+00 | 0 | 0 |
| amino acids, aromatic | 3 | 1.00E+00 | 0.00E+00 | 0 | 0 |
| amino acids, basic | 3 | 1.00E+00 | 0.00E+00 | 1 | 0 |
| amino acids, branched-chain | 3 | 1.00E+00 | 0.00E+00 | 1 | 1 |
| amino acids, cyclic | 4 | 1.00E+00 | 0.00E+00 | 1 | -1 |
| amino acids, sulfur | 5 | 1.00E+00 | 0.00E+00 | 0 | -1 |
| benzene derivatives | 3 | 1.00E+00 | 0.00E+00 | 1 | 0 |
| biogenic polyamines | 4 | 1.00E+00 | 0.00E+00 | 1 | -1 |
| cholestenes | 4 | 1.00E+00 | 0.00E+00 | 2 | 1 |
| dicarboxylic acids | 5 | 7.30E-03 | 4.92E+00 | 0 | -1 |
| dipeptides | 5 | 1.00E+00 | 0.00E+00 | 2 | 0 |
| disaccharides | 6 | 2.20E-01 | 1.51E+00 | 0 | -1 |
| ethanolamines | 3 | 1.00E+00 | 0.00E+00 | 0 | 0 |
| glutarates | 3 | 1.00E+00 | 0.00E+00 | 0 | 0 |
| glyceric acids | 3 | 1.00E+00 | 0.00E+00 | 0 | 0 |
| glycerides | 3 | 1.00E+00 | 0.00E+00 | 0 | 0 |
| hexosephosphates | 6 | 1.00E+00 | 0.00E+00 | 1 | 0 |
| hexoses | 3 | 1.00E+00 | 0.00E+00 | 0 | -1 |
| pentoses | 3 | 1.00E+00 | 0.00E+00 | 2 | 0 |
| purine nucleosides | 7 | 4.20E-02 | 3.17E+00 | 1 | 1 |
| purinones | 3 | 1.00E+00 | 0.00E+00 | 0 | -1 |
| pyrimidine nucleosides | 3 | 1.00E+00 | 0.00E+00 | 0 | 0 |
| pyrimidinones | 3 | 1.00E+00 | 0.00E+00 | 4 | 0 |
| saturated fatty acids | 14 | 1.80E-01 | 1.71E+00 | 4 | 0.1 |
| sugar alcohols | 9 | 2.70E-02 | 3.61E+00 | 1 | 0.1 |
| unsaturated fatty acids | 6 | 1.00E+00 | 0.00E+00 | 4 | -1 |

**Supplemental Table XIII:** ChemRICH cluster information for 2.0MOI_36HPI.

| Cluster name | Cluster size | P-value | Transformed P-value | Altered metabolites | Up/down ratio |
| --- | --- | --- | --- | --- | --- |
| adipates | 3 | 1.00E+00 | 0.00E+00 | 1 | -1.0 |
| amino acids | 9 | 3.90E-03 | 5.55E+00 | 5 | -0.8 |
| amino acids, acidic | 3 | 1.00E+00 | 0.00E+00 | 1 | -1.0 |
| amino acids, aromatic | 3 | 1.00E+00 | 0.00E+00 | 0 | 0 |
| amino acids, basic | 3 | 1.00E+00 | 0.00E+00 | 0 | 0 |
| amino acids, branched-chain | 3 | 1.00E+00 | 0.00E+00 | 0 | 0 |
| amino acids, cyclic | 4 | 1.00E+00 | 0.00E+00 | 1 | -1.0 |
| amino acids, sulfur | 5 | 1.00E+00 | 0.00E+00 | 2 | -1.0 |
| benzene derivatives | 3 | 1.00E+00 | 0.00E+00 | 0 | 0 |
| biogenic polyamines | 4 | 1.10E-01 | 2.21E+00 | 2 | -1.0 |
| cholestenes | 4 | 1.00E+00 | 0.00E+00 | 0 | 0 |
| dicarboxylic acids | 5 | 7.30E-03 | 4.92E+00 | 2 | -1.0 |
| dipeptides | 5 | 1.00E+00 | 0.00E+00 | 0 | 0 |
| disaccharides | 6 | 7.80E-02 | 2.55E+00 | 2 | -1.0 |
| ethanolamines | 3 | 1.00E+00 | 0.00E+00 | 0 | 0 |
| glutarates | 3 | 1.00E+00 | 0.00E+00 | 0 | 0 |
| glyceric acids | 3 | 1.00E+00 | 0.00E+00 | 0 | 0 |
| glycerides | 3 | 1.00E+00 | 0.00E+00 | 1 | -1.0 |
| hexosephosphates | 6 | 1.00E+00 | 0.00E+00 | 0 | 0 |
| hexoses | 3 | 1.00E+00 | 0.00E+00 | 1 | -1.0 |
| pentoses | 3 | 1.00E+00 | 0.00E+00 | 0 | 0 |
| purine nucleosides | 7 | 1.00E+00 | 0.00E+00 | 1 | 1 |
| purinones | 3 | 1.00E+00 | 0.00E+00 | 1 | -1.0 |
| pyrimidine nucleosides | 3 | 1.00E+00 | 0.00E+00 | 0 | 0 |
| pyrimidinones | 3 | 1.00E+00 | 0.00E+00 | 0 | 0 |
| saturated fatty acids | 14 | 7.00E-02 | 2.66E+00 | 2 | 0.1 |
| sugar alcohols | 9 | 4.50E-02 | 3.10E+00 | 2 | 0.1 |
| unsaturated fatty acids | 6 | 5.30E-03 | 5.24E+00 | 3 | -1.0 |

**Supplemental Figure 1:**
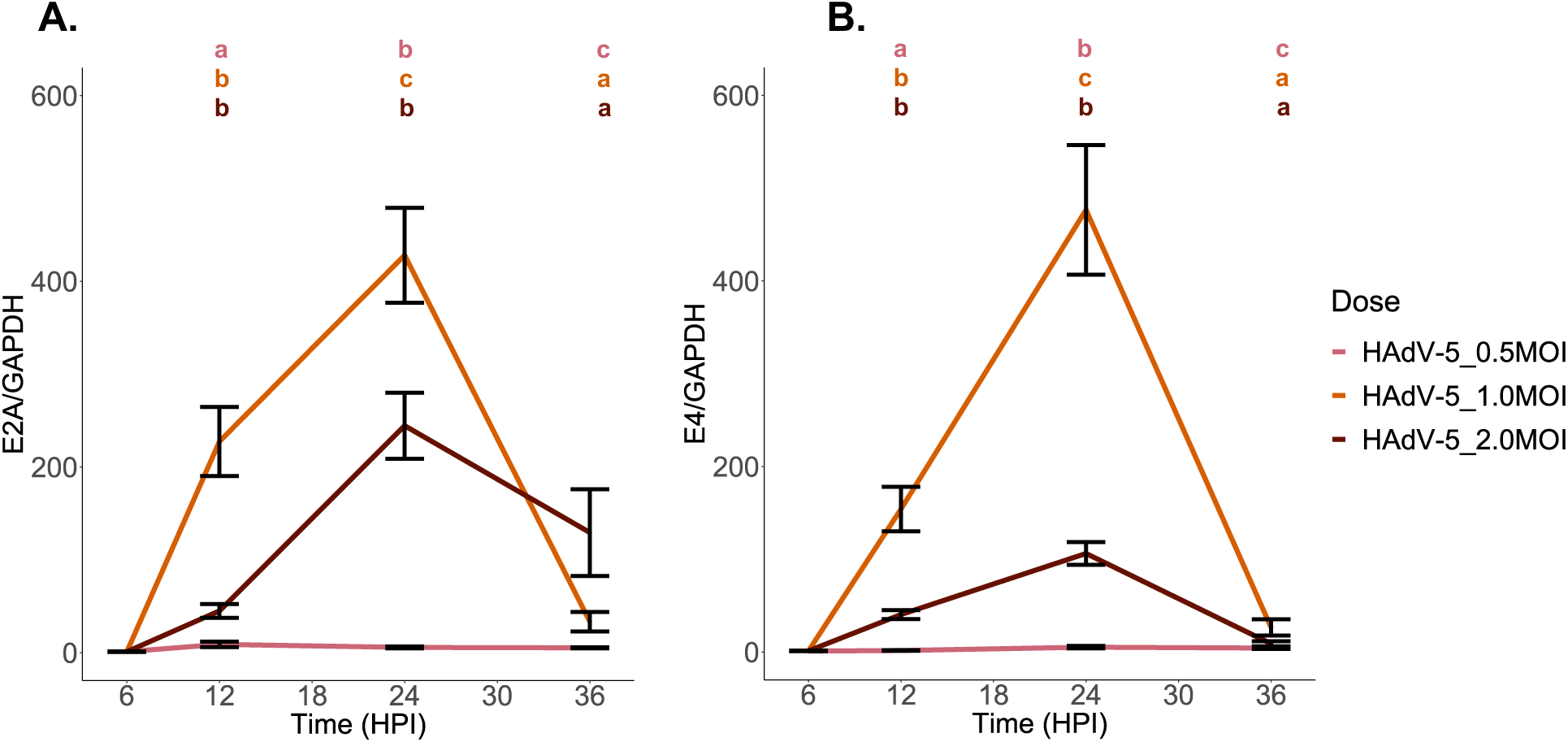
RT-qPCR data confirming viral infection at all time points in this study. **A**. Relative gene expression results for Pan E2A gene for 0.5MOI, 1.0MOI, and 2.0MOI dosages across time. **B.** Relative gene expression results for Pan E4 gene for 0.5MOI, 1.0MOI, and 2.0MOI dosages across time. Experiments were performed in sextuplicate. Data are presented as the standard error of the mean (SEM). **a** = P < 0.05, **b** = P < 0.01, **c** = P < 0.001, **ns** = not significant.

**Supplemental Figure 2:**
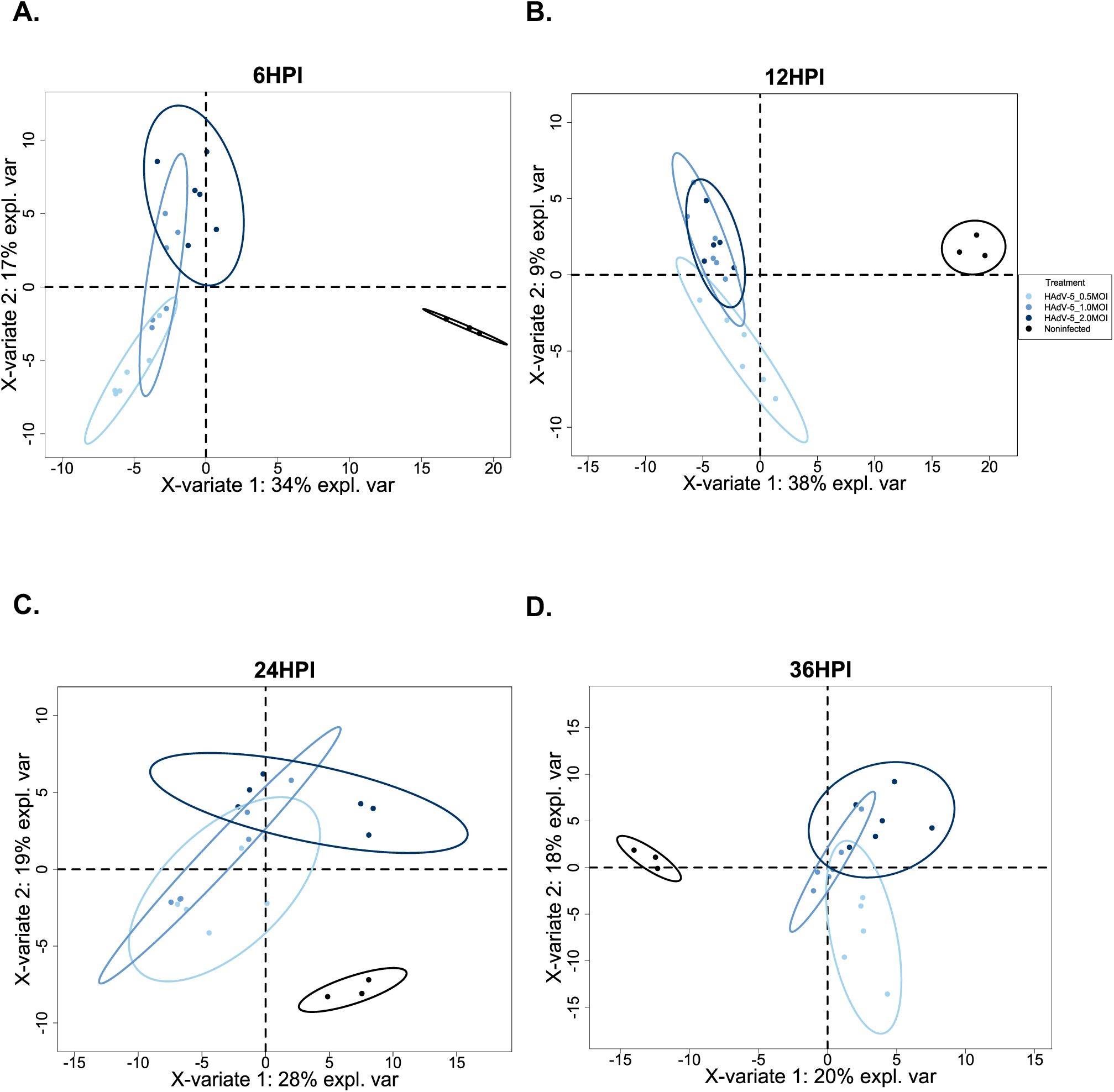
Partial least squares-discriminant analysis (PLS-DA) revealed separation between the control and experimental conditions. **A**. PLS-DA for 6HPI comparisons. **B.** PLS-DA for 12HPI comparisons. **C.** PLS-DA for 24 HPI comparisons. **D.** PLS-DA comparison for 36HPI comparisons.

**Supplemental Figure 3:**
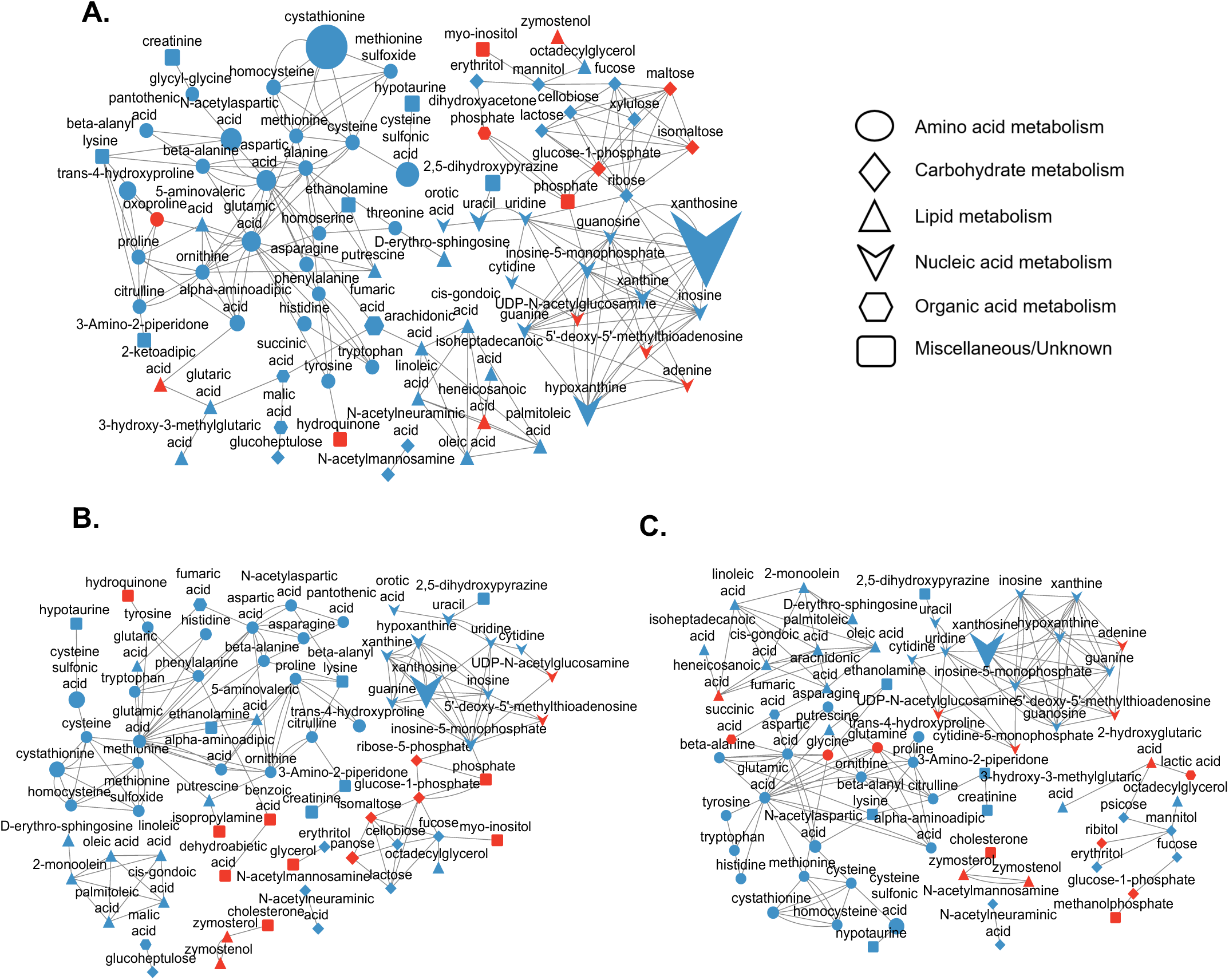
Network analyses at 6 HPI show early perturbation of individual metabolites across multiple categories. **A**. Network map analyzing the biochemical relationships between metabolites significantly altered in the 0.5MOI/noninfected comparison. **B.** Network map analyzing the biochemical relationship between metabolites significantly altered in the 1.0MOI/noninfected comparison. **C.** Network map analyzing the biochemical relationships between metabolites significantly altered in the 2.0MOI/noninfected comparison. The size of each shape corresponds to the fold change value calculated using MetaMapp. Blue shapes represent downregulated metabolites, whereas red shapes indicate upregulated metabolites.

**Supplemental Figure 4:**
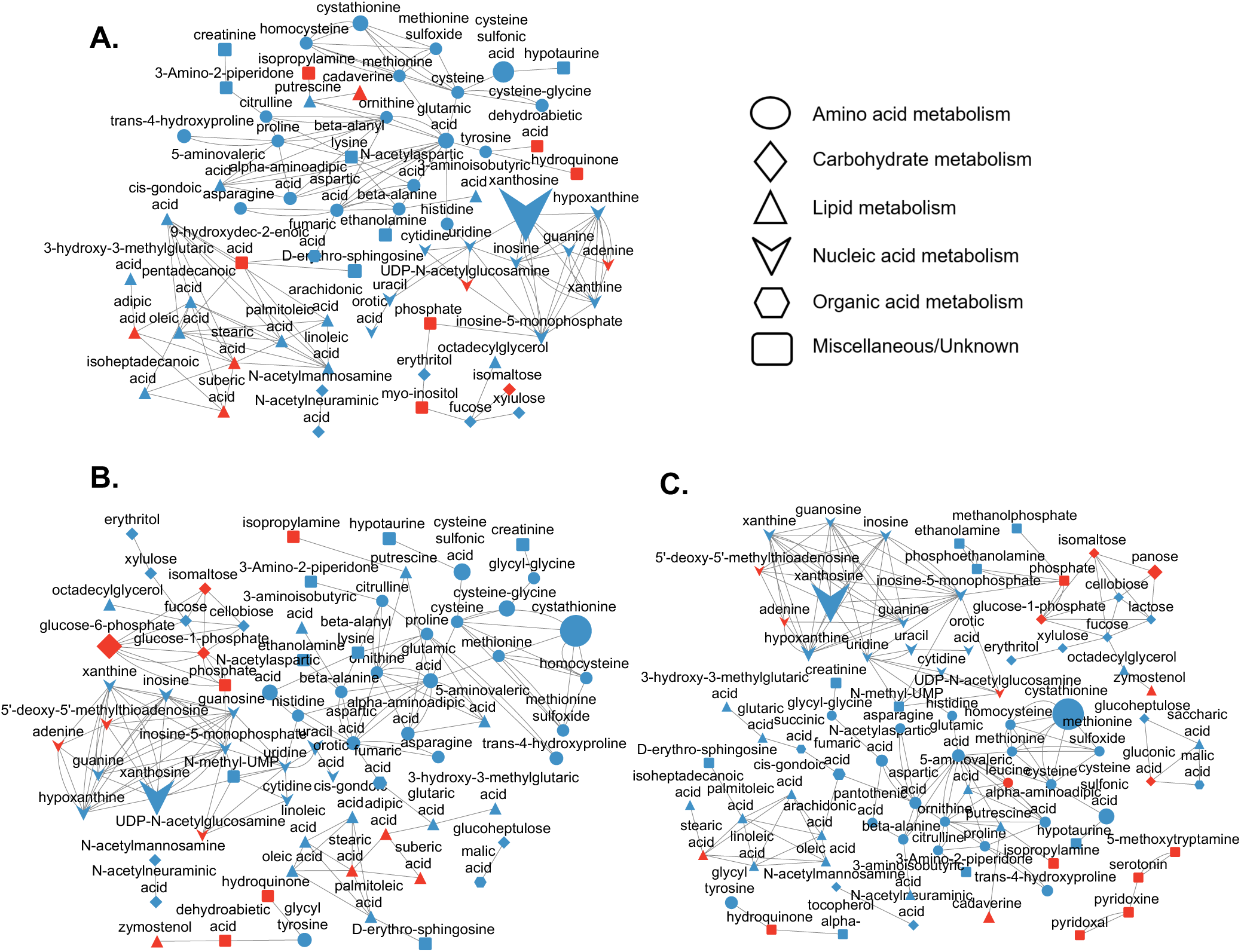
Network analyses reveal continued metabolic perturbations at 12 HPI. **A** Network map analyzing the biochemical relationships between metabolites significantly altered in the 0.5MOI/noninfected comparison. **B** Network map analyzing the biochemical relationship between metabolites significantly altered in the 1.0MOI/noninfected comparison. **C** Network map analyzing the biochemical relationships between metabolites significantly altered in the 2.0MOI/noninfected comparison. The size of each shape corresponds to the fold change value calculated using MetaMapp. Blue shapes represent downregulated metabolites, whereas red shapes indicate upregulated metabolites.

**Supplemental Figure 5:**
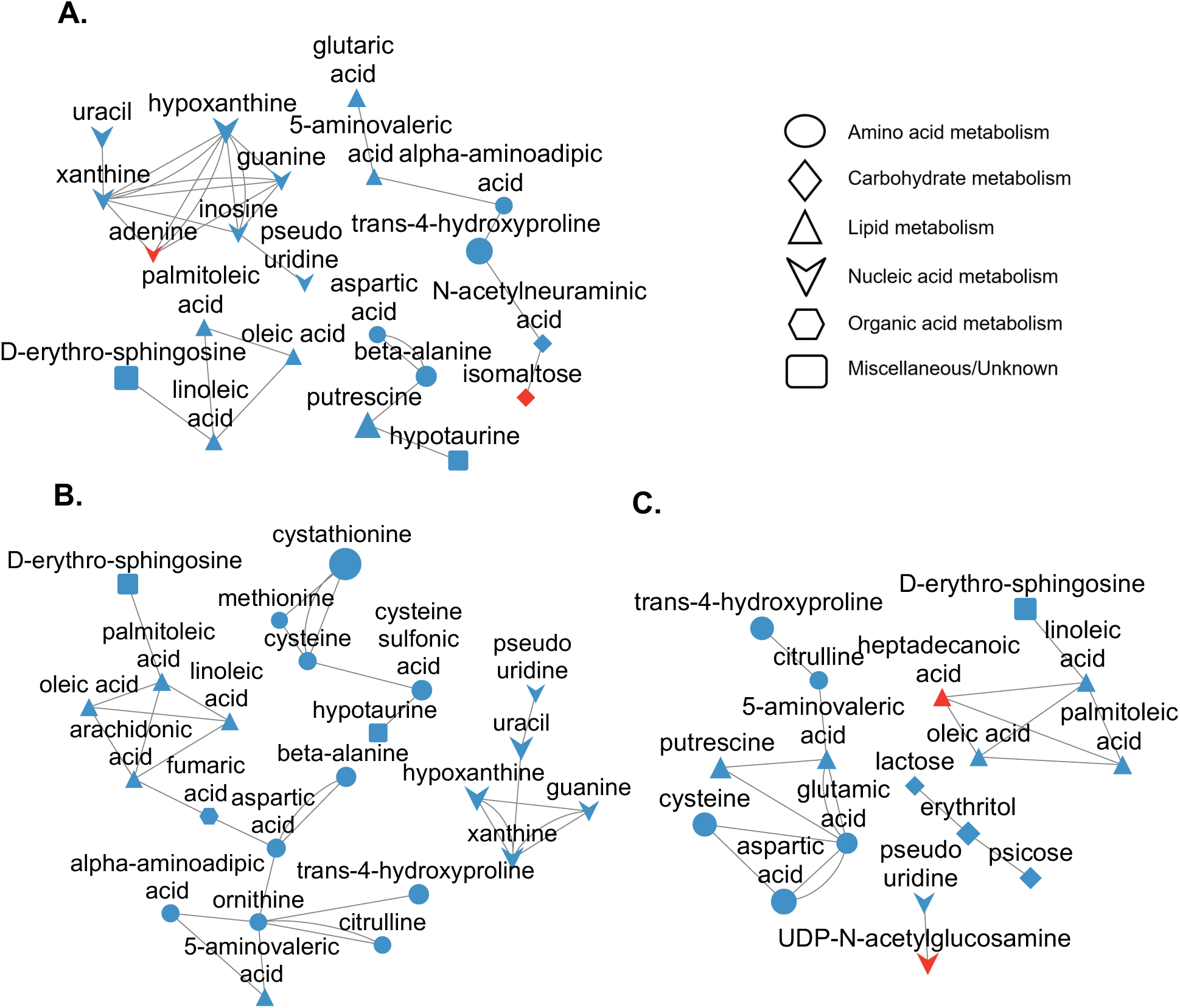
Network analyses at 24HPI reveal a marked reduction in metabolite involvement across amino acid, lipid, and nucleic acid metabolism. **A**. Network map analyzing the biochemical relationships between metabolites significantly altered in the 0.5MOI/noninfected comparison. **B.** Network map analyzing the biochemical relationship between metabolites significantly altered in the 1.0MOI/noninfected comparison. **C.** Network map analyzing the biochemical relationships between metabolites significantly altered in the 2.0MOI/noninfected comparison. The size of each shape corresponds to the fold change value calculated using MetaMapp. Blue shapes represent downregulated metabolites, whereas red shapes indicate upregulated metabolites.

**Supplemental Figure 6:**
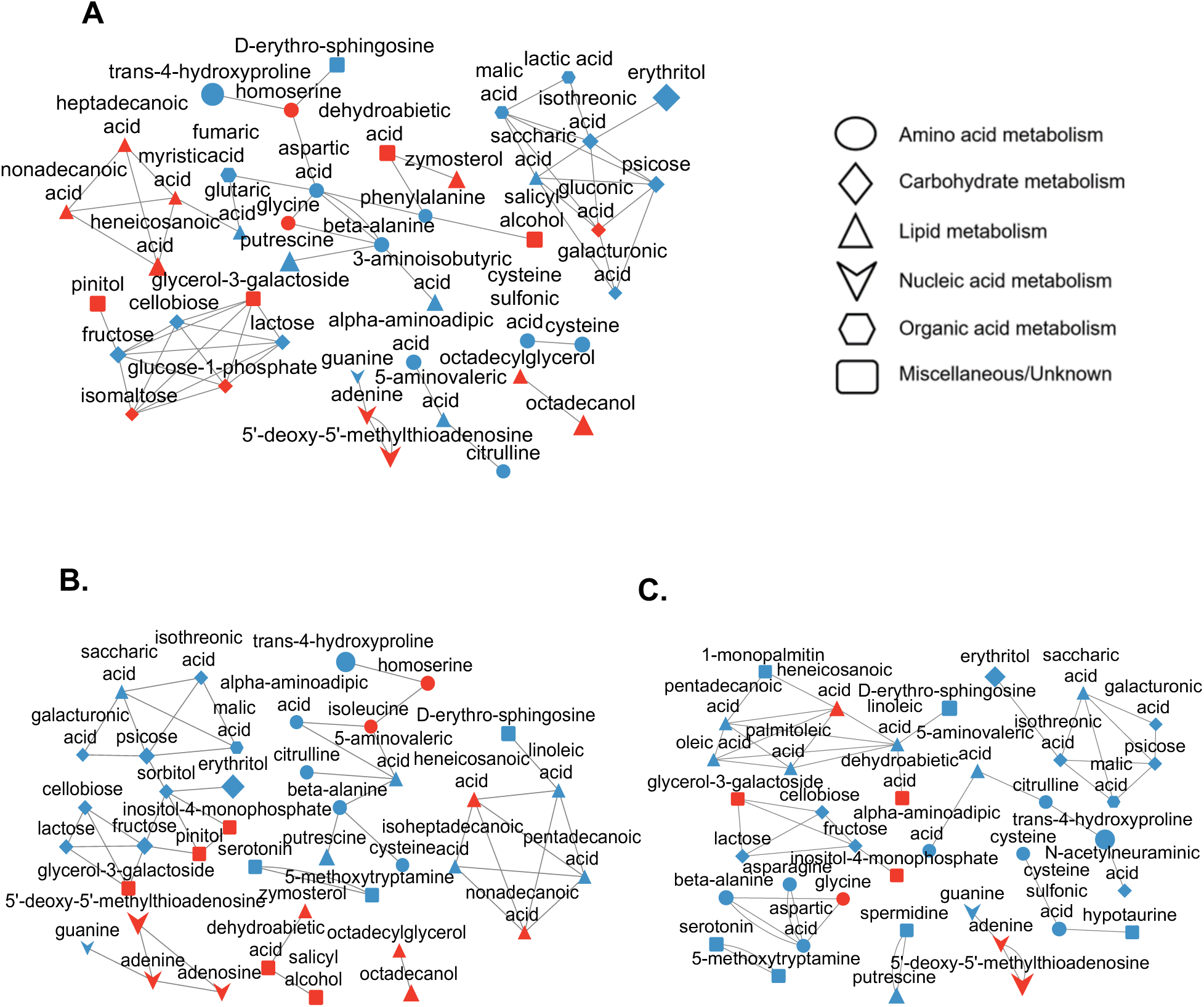
Network analyses at 36HPI yield increases in metabolite shifts across treated/untreated comparisons. **A**. Network map analyzing the biochemical relationships between metabolites significantly altered in the 0.5MOI/noninfected comparison. **B.** Network map analyzing the biochemical relationship between metabolites significantly altered in the 1.0MOI/noninfected comparison. **C.** Network map analyzing the biochemical relationships between metabolites significantly altered in the 2.0MOI/noninfected comparison. The size of each shape corresponds to the fold change value calculated using MetaMapp. Blue shapes represent downregulated metabolites, whereas red shapes indicate upregulated metabolites.

